# Sincast: a computational framework to predict cell identities in single cell transcriptomes using bulk atlases as references

**DOI:** 10.1101/2021.11.07.467660

**Authors:** Yidi Deng, Jarny Choi, Kim-Anh Lê Cao

**Author notes:** indicates equal contribution (Jarny Choi, Kim-Anh Lê Cao).

## Abstract

Characterizing the molecular identity of a cell is an essential step in single cell RNA-sequencing (scRNA-seq) data analysis. Numerous tools exist for predicting cell identity using single cell reference atlases. However, many challenges remain, including correcting for inherent batch effects between reference and query data and insufficient phenotype data from the reference. One solution is to project single cell data onto established bulk reference atlases to leverage their rich phenotype information.

Sincast is a computational framework to query scRNA-seq data based on bulk reference atlases. Prior to projection, single cell data are transformed to be directly comparable to bulk data, either with pseudo-bulk aggregation or graph-based imputation to address sparse single cell expression profiles. Sincast avoids batch effect correction, and cell identity is predicted along a continuum to highlight new cell states not found in the reference atlas.

In several case study scenarios, we show that Sincast projects single cells into the correct biological niches in the expression space of the bulk reference atlas. We demonstrate the effectiveness of our imputation approach that was specifically developed for querying scRNA-seq data based on bulk reference atlases. We show that Sincast is an efficient and powerful tool for single cell profiling that will facilitate downstream analysis of scRNA-seq data.

## 1 Introduction

Single cell RNA sequencing (scRNA-seq) allows for the study of cell-specific variations in transcriptional states at an unprecedented resolution. One essential step in scRNA-seq data analysis is to characterize cell molecular identity, either *de novo* or with existing vocabularies of known cell types or states. Numerous computational tools have been developed for predicting cell identity using other single cell atlases as references (Andreatta et al., 2021; Clarke et al., 2021). However many challenges remain, including integrating atlases from independent studies to build comprehensive atlases that are generalizable, annotating reference cells accurately and tuning the parameters of these prediction tools appropriately (Zhao et al., 2020). Furthermore, the reference and query data effectively represent separate batches. Correcting for batch effects is required before direct comparisons can be made. Using data integration to address this issue is difficult from both a statistical and data analysis perspective (Argelaguet et al., 2021; Luecken et al., 2020). During the reference-query integration task, biological and batch effects are confounded, resulting in the potential removal of large amount of biological variation that is considered as batch variation.

In light of these challenges, bulk sequencing data represent a valuable resource for building reference atlases, as the samples can be of high quality, well replicated and well annotated as their phenotype is known (Angel et al., 2020; Chandra et al., 2021; Choi et al., 2019; Davis et al., 2018; Lizio et al., 2015; Mabbott et al., 2013; Rajab et al., 2021). However, using bulk atlases for single cell identity has mostly been overlooked. Instead, some studies have proposed to analyse bulk data using scRNA-seq data as a reference. For example, many deconvolution methods have been developed to estimate bulk sample cellular composition based on scRNA-seq (Cobos et al., 2020; Kuksin et al., 2021). Only a few approaches have attempted to decipher cellular identity of scRNA-seq by leveraging bulk data. SingleR annotates query cells using labels of bulk reference samples that are matched to each cell according to Spearman correlation (Aran et al., 2019). Capybara predicts continuous cell identity by regressing each query cell expression profile on a bulk reference with restricted linear regression (Kong et al., 2020). SCRABBLE imputes scRNA-seq under the constraint that the averaged expression of imputed single cells is consistent with a given bulk reference (Peng et al., 2019). These methods remain challenged by large technical differences between scRNA-seq and bulk data, in particular library size and zero composition (Sarkar and Stephens, 2021). Roels et al. (2020) addressed this challenge by down sampling reads in reference bulk data prior to data integration with scRNA-seq query using the approach from Seurat (Hao et al., 2021).

We propose a computational framework, Sincast, to query scRNA-seq data against bulk transcriptional reference atlases. Our framework avoids reliance on data integration to address technical differences across batches (Figure 1). Our bulk transcriptional reference atlases can include both microarray and RNA-seq data, and are built based on Principal Component Analysis (PCA), as we previously proposed in Angel et al. 2020. Single cell RNA-seq query data are projected onto the low-dimensional expression space spanned by the atlas principal components. The location of the query cells on the atlas allows the identification of similarities with well annotated bulk reference samples. Prediction of cell identity is based on an improved Capybara score (Kong et al., 2020). Most importantly, the core challenge of the structural differences between the reference and the query is addressed with two independent approaches, depending on the data structure of the query. We propose to either aggregate single cells to create pseudo-bulk samples, mimicking structure and variation of bulk samples, or to zero-impute single cell data as sparsity is a major data characteristic that deviates single cell from bulk data. Our proposed rank transformation of the query and the atlas profiles independently also avoid the need of batch effect correction (Angel et al., 2020). On five case studies (each query being projected on a relevant reference atlas), we demonstrate that we can robustly map single cells into correct biological niches of bulk atlases with a high concordance with the biology described in the original query study. We also show that new and more continuum cell states can be predicted through Sincast projection, and that key regulators of differentiation and pseudo-time trajectories can be obtained without the need for complex algorithms by leveraging the principal components of the reference atlas.

**Figure 1.**
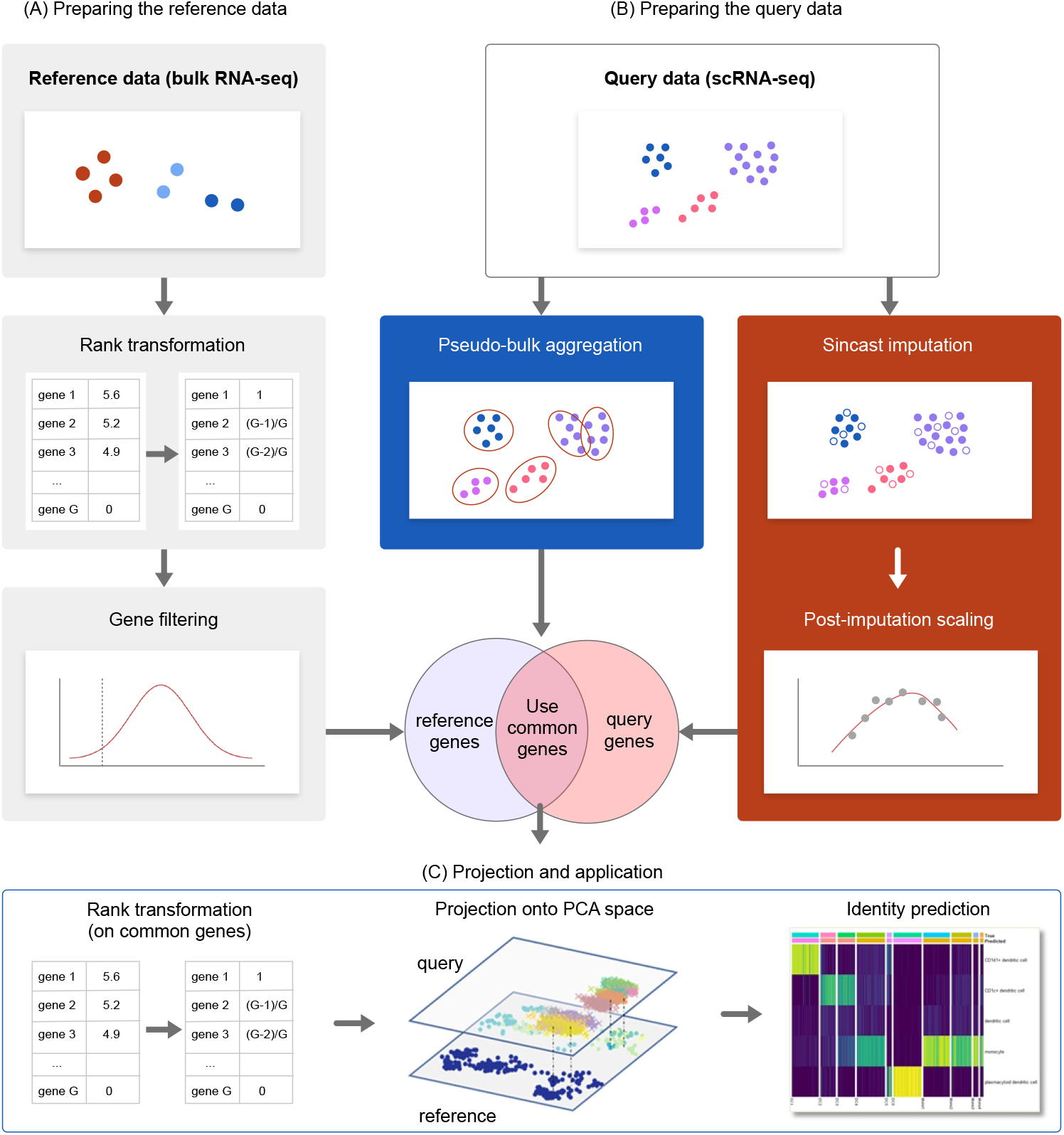
An overview of Sincast framework for projecting query scRNA-seq data onto reference bulk atlas. The differences in zero composition and scale between bulk and scRNA-seq data constitute major challenges to capture biologically relevant variation in the single cells, which Sincast addresses without data integration. **(A)** The reference bulk data are rank transformed, as proposed by Angel et al. 2020 and additional gene filtering based on Hellinger Distance is applied to retain the most important genes discriminating cell types. **(B)** For the query single cell data, Sincast either aggregates single cells by pooling the expression profiles of cells to create pseudo bulk samples, or zero imputes the data by inferring unobserved expressions in a cell from the other cells in the query, followed by robust data normalization. The overlapping genes are then rank transformed for **(C)** projection, which consists in aligning both query and reference. Principal Component Analysis is performed on the reference data to construct a low dimensional expression space (atlas). Projection of the query is performed by calculating the query principal component scores learnt from the reference, and projection is further improved by diffusion map. Cell identity prediction based on the neighboring reference samples on the atlas is performed with a modified Capybara score (Kong et al., 2020).

## 2 Results

### 2.1 Projecting data after pseudo-bulk aggregation is a simple and effective way to reveal cell identity

We found that projecting single cell data onto a bulk reference without any transformation performed poorly. The projected cells tended to cluster together, away from all other reference clusters rather than co-localised to the same biologically matching bulk samples (Figure 2A). This result was not surprising due to the large difference in data structure between single cell and bulk - in particular the zero inflated nature of single cell data.

**Figure 2.**
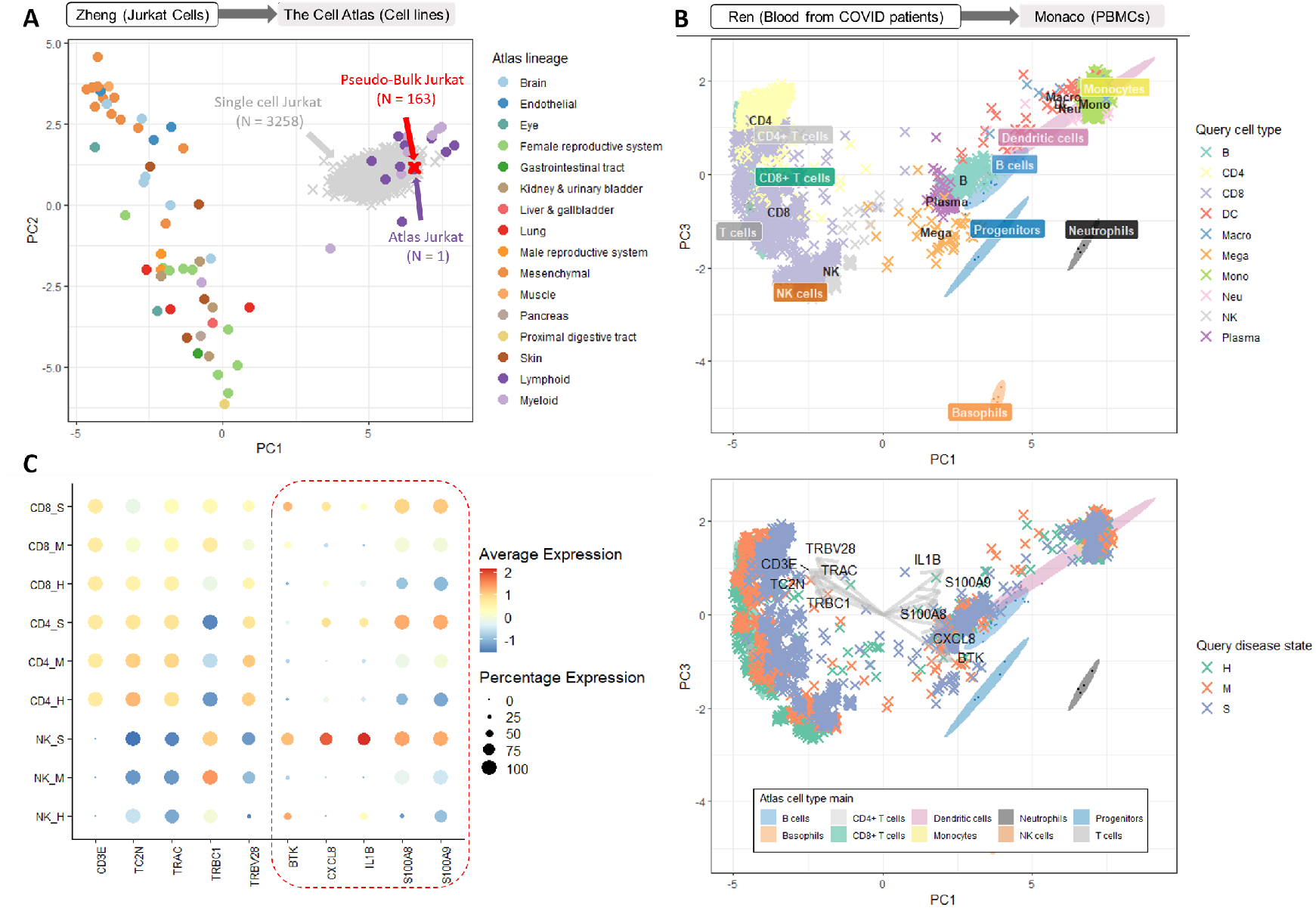
**(A)** We projected single cell data from Zheng et al. (2017), representing Jurkat cells profiled using 10x Genomics, onto The Cell Atlas (Thul et al., 2017) representing bulk RNA-Seq profiles of cell lines. Projection without any transformation resulted in the cells (in grey) being identified as lymphoid cells. After pseudo-bulk aggregation the cells (in red) projected closest to the Jurkat cells in the reference. **(B)** We projected immune cells from COVID-19 infected donors as well as healthy controls (Ren et al., 2021) onto bulk RNA-seq atlas of immune cells (Monaco et al., 2019) after pseudo-bulk aggregation. The cells were projected accurately onto the corresponding cell types of the reference (top). When we coloured the same projected cells by disease state (bottom), we observed a clear shift in the identities of lymphoid cells according to disease severity (H: healthy, M: medium, S: severe). The arrows represent the top five positive and top five negative loading important genes that define PC1. **(C)** Dot plot showing the expression of the top loading genes as described in (B), highlighting an increase in the expression of each of the positive loading genes with disease severity.

Pseudo-bulk aggregation is a straightforward way to make single cell data compatible for projection onto bulk. Aggregation is done by sampling cells of the same cluster with replacement and adding up their expression profiles. This approach is simple to implement and also conforms to our biological understanding that bulk expression represents pooled single cell expression. We illustrate the usefulness of this approach through two case studies, where the query and reference data contain biologically matching cell types.

#### Case study 1: projecting Jurkat cells onto The Cell Atlas shows pseudobulk aggregation can classify cells accurately

The reference atlas from The Cell Atlas (Thul et al., 2017) consists of bulk RNA-seq data from a comprehensive range of cell lines. The query data from Zheng et al. (2017) contain Jurkat T cell line from 10x Genomics (3,2058 cells, see also Table 1). Principal Component Analysis (PCA) of the reference data showed a strong separation of blood cells from the other cell types along the first principal component (PC1, Figure 2A). However, even though the non aggregated single cell data were projected onto the blood cell area of the PCA space, classifying them as one of the nearby cell types was difficult. Pseudo-bulk aggregation was more successful, as all aggregated cells were projected very closely to the Jurkat cell of the reference.

**Table 1.** Summary of the case studies, including the reference data on which we built the atlases and their number of samples, the query data for the corresponding reference atlases, and their numbers of cells, the number of discriminant genes selected for the reference atlas and their overlap between the reference and query prior to projection.

| Reference Data | Reference cells | Query Data | Query Cells | Genes Selected and Overlap | Used in |
| --- | --- | --- | --- | --- | --- |
| Thul et al. (2017)<br>RNA-seq of 69 Cell lines | N = 69 | Zheng et al. (2017)<br>Single cell Jurkat T (10x v1) | N = 3,258 | 3,000/1,531 | Section 2.1<br>Figure 2A |
| Schmiedel et al. (2018)<br>Gene expression data from the DICE project. | N = 1,561 | Zheng et al. (2017)<br>Single cell Jurkat T (10x v1) | N = 3,258 | 2,000/1,556 | Section 5.8<br>Figure S1 |
|  |  | Lizio et al. (2015)<br>Bulk Jurkat T (Fantom5) | N = 1 |  |  |
|  |  | Davis et al. (2018)<br>Bulk Jurkat T (ENCODE, identity: ENCSR000BXX) | N = 1 |  |  |
| Monaco et al. (2019)<br>Molecular characterization of 29 immune cells within peripheral blood mononuclear cell. | N = 114 | Ren et al. (2021)<br>Human immune response to COVID19 infection | N = 49,900 | 1,000/937 | Section 2.1, 5.9<br>Figure 2B,C<br>S2, S3, S6 |
| Rajab et al. (2021)<br>An integrated myeloid atlas | N = 901 | Bian et al. (2020)<br>Deciphering human embryonic macrophage development | N = 1,231 | 2,000/1,952 | Section 2.2<br>Figure 4<br>S4, S10, S12 |
| Monocyte and DC subset of Rajab et al. (2021) | N = 500 | Villani et al. (2017)<br>Human dendritic cell and monocyte subsets | N = 1,078 | 500/416 | Section 2.2, 5.10<br>Figure 3<br>S5, S11, S13, S15 |

#### Case study 2: querying COVID-19 case-control study data onto an immune cell atlas shows pseudo-bulk aggregation can highlight shifts in cell identity

The reference data from Monaco et al. (2019) consist of 29 immune cells sorted from peripheral blood mononuclear cells. The query cells were from Ren et al. (2021), describing immune cells profiled on both healthy and COVID-19 infected donors. We selected nine donors from the same batch, in different disease stages of healthy, moderate and severe, to aggregate and project (see also Table 1).

Figure 2B illustrated the pseudo-bulk aggregated projection coloured by cell type only (See Supplementary Figure S2 for the projection colored by atlas and query cell type). We observed a high concordance between query and reference cell types. Next, we coloured the projected cells according to disease stage on the same plot. This projection illustrated that the T and the NK cell populations of COVID patients had identity shifts towards the positive direction of PC1 of the reference compared to the healthy controls (Figure 2B). We found that inflammatory markers such as BTK, CXCL8, IL1B, S100A8/9, were among the top 20 genes with the highest PC1 loadings (i.e. important genes that drive linear separation of samples on PC1). The shifts of cell population indicated an up-regulation of these inflammatory signatures in COVID patients according to disease severity (Figure 2C). This finding was consistent with Ren et al. (2021), who claimed that hyper-inflammatory cell subtypes defined by the systematical up-regulation of these inflammatory signatures were one of the major causes of cytokine storm in severe COVID patients.

In the myeloid compartment of the projection, the shift in the projected monocytes of COVID patients compared to the healthy controls was difficult to visualize. Thus we applied our improved Capybara cell score to the projected cells (Kong et al., 2020) to quantify the projection more rigorously. Our predicted score revealed that non-classical monocytes (CD14-CD16+) in COVID patients acquired an intermediate monocyte (CD14+ CD16+) identity (Supplementary Figure S3), providing potential explanation on the reported increase of intermediate monocytes in the PBMCs of COVID patients (Zhang et al., 2020; Zhou et al., 2020).

This case study showed that pseudo-bulk aggregation can work beyond simply benchmarking cells when there is high concordance between the query and reference cell types, and can reveal more intermediate cell types. It also illustrated how a projection method can rapidly generate biological insight, without the need to perform differential expression analysis separately for example. Indeed, the reference atlas already contained key genes which defined the principal components in the PCA space. Batch correction was not necessary when projecting, a feature from Sincast that provides a large advantage when the query data contain large batch effects. It is possible to extend this idea even further by using the reference as a background on which multiple query data can be compared to each other without batch correction (see Supplementary Material 5.8, Figure S1).

#### Limitations of pseudo-bulk aggregation

Case study 2 (Figure 2B) highlighted some ‘mismatched’ cell projections near the centre of the PCA space, illustrating an inherent limitation of pseudo-bulk aggregation when the query cluster is highly sparse. Aggregation requires a sufficient number of bootstrap sampling from each cluster to overcome zero-inflation problem. Thus, a cluster composed of only a few cells poses a problem as the pooled gene counts may still be zero-inflated.

We defined sparsity in this context as the percentage of zeros present in a pseudo-bulk aggregated cluster. We assessed whether a sparsity threshold could indicate the appropriateness of pseudo-bulk aggregation, depending on the study and cell types. We down sampled the atlas samples to simulate sparse samples to project. The threshold was defined at the point where matched cell identities of sparse samples diverged (Supplemental Material 5.9). For example, case study 2 showed that any cluster with sparsity greater than 15 percent led to poor projection (Supplemental Figure S6, and Figures S7, S8, S9, for other case studies).

In addition, we found that pseudo-bulk aggregation was limited when the query data did not contain very distinct clusters, or when intra-cluster variance was large. Single cell data are often described as a high resolution version of bulk data, where continuum cell states may be more readily observed. Cluster assignments may be ambiguous or dependent largely on the choice of parameters. One way to address this issue with pseudo-bulk aggregation is to use an arbitrary number of sub-clusters to aggregate, and vary this number according to some metric, but interpretation of the projections will be difficult when dealing with such a random array of sub-clusters.

### 2.2 Data imputation prior to projection reveals complex single cell biology

When the query data contain clusters with high sparsity or represent a more continuum of cell states rather than distinct states, data imputation offers an alternative to pseudo-bulk aggregation. However, we show that existing imputation methods created inaccurate projections, due to over smoothing of the query data prior to projection, resulting in over-shrinking the variance. Our imputation method builds on MAGIC (Van Dijk et al., 2018) to project single cell data onto bulk reference. We compare our method against existing imputation methods in two case studies and illustrate how imputation followed by projection can reveal new cell states.

#### Case study 3: Existing scRNA-seq imputation methods show limitations when used to project onto bulk reference data

We considered the reference data from Rajab et al. (2021), where we previously integrated 44 microarray and bulk RNA-seq datasets to create an atlas of myeloid cells. The query data from Bian et al. (2020) contain myeloid cells derived from human embryos (see also Table 1). Three existing single cell imputation methods were compared with their default parameters: MAGIC (Van Dijk et al., 2018), knn-smothing (Wagner et al., 2017), and SAVER (Huang et al., 2018). These methods chosen as they were the top three performers in the review of imputaton methods by Hou et al. (2020).

We found that the projection of imputed single cell data onto the reference differed greatly depending on the imputation method, reflecting the assumptions and characteristics of each method (Supplementary Figure S10). In this case study, cells imputed by MAGIC were connected to form smooth cellular trajectories with restricted local variance. Cells imputed by knn-smoothing were more scattered than MAGIC, as a result of iterative data aggregation during imputation. Cells imputed by SAVER, a model-based method that predicts the expression profile of each cell by regressing on the rest of the cells, were not shrunk locally relative to the global scale of the query data. The projection visualisation can be used as preliminary benchmark to assess the relevance of these methods in this context.

To illustrate how these differences translated to specific projection results, we focused on the embryonic macrophages Mac 1 and Mac 4 in the query data. Bian et al. (2020) noted that these describe distinct cell identities, where Mac 1 cells were mainly found in the yolk sac at Carnegie Stage 11, while Mac 4 cells were predominantly located in the head representing developing microglia. Only the projection made after MAGIC or Sincast imputation showed these cell types as distinct clusters (Figure S12).

#### Case study 4: Sincast imputation produces more accurate projections onto bulk reference data

We next evaluated the performance of Sincast imputation against these three imputation methods. This case study used the query data from Villani et al. (2017), which contain 6 dendritic cell (DC) sub-populations, FACS sorted and profiled using Smart-seq2 (Picelli et al., 2013). For the bulk reference, we chose a pseudo-bulk aggregated version of the query data itself and used the accompanying annotation as ground truth in the evaluation (see also Table 1). We also calculated median silhouette index (MSI) and adjusted rand index (ARI) on the query projection to evaluate the accuracy of the results. MSI and ARI measure how well each cell’s cluster membership is preserved before and after imputation.

While all imputation methods improved cell type classification compared to raw data projection, Sincast imputed data performed best in terms of ARI and the second best in terms of MSI (Figure 3A). Each of the DC clusters of Villani et al. (2017) projected onto their matched reference cell types after Sincast imputation. We then evaluated the robustness of Sincast regarding its imputation tuning parameters on the same atlas, compared to MAGIC (see Method Section 4.5). We only imputed and projected ten DC6 and 285 Mono1/Mono2 cells of the query (see Supplementary Material 5.10). We intentionally imputed each cell based on its 15 nearest neighbors (i.e a value larger than the actual DC6 population), and varied the diffusion time parameter *t* for both MAGIC and Sincast before projection. With MAGIC, higher values of *t* resulted in the DC6 cluster from the query data projecting further from the reference DC6 cell cluster, towards monocytes. We did not observe such effect with Sincast (Figure 3B).

**Figure 3.**
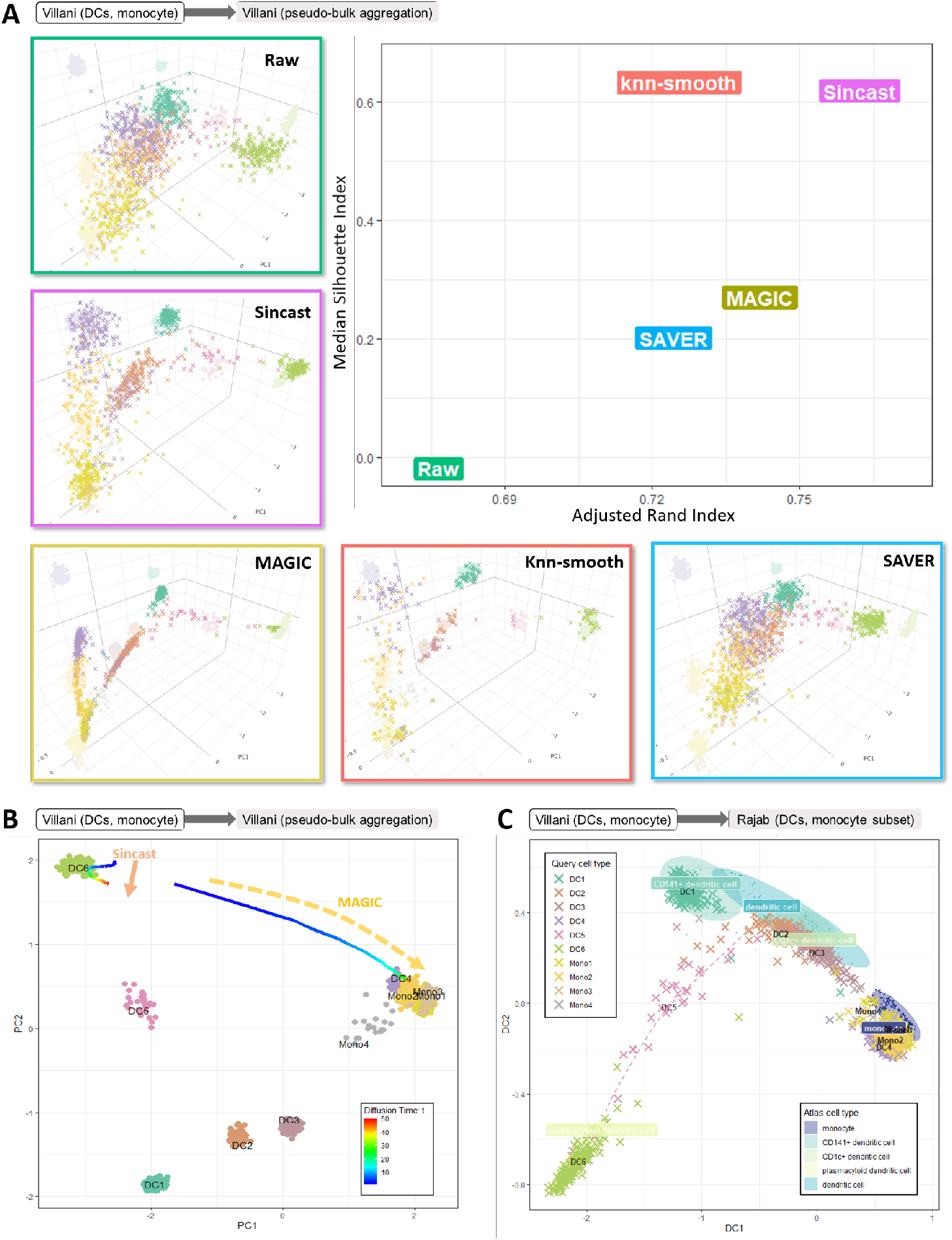
**(A)** We projected the DC cells from Villani et al. (2017) onto a pseudo-bulk version of the same data to evaluate the performance of popular imputation methods in the context of projection. Measures of accuracy such as adjusted rand index and median silhouette showed Sincast performed best. **(B)** To assess impact of imputation tuning parameters on the projection results, we imputed then projected the subset of DC6, Mono1 and Mono2 cells from Villani et al. (2017) onto the Villani pseudo-bulk atlas while varying the diffusion time parameter *t* for MAGIC and Sincast. The line shows the centroids of projected points according to *t* values. The DC6 population after MAGIC imputation was wrongly assigned monocyte identity when *t* increased, unlike Sincast imputation which preserved the DC6 identity. **(C)** By reconstructing the PCA projection landscape with diffusion map, Sincast imputed version of Villani et al. (2017) projected the cells accurately onto the bulk DC and monocyte subset of Rajab et al. (2021). The projection also highlighted the newly discovered DC5 population as a continuum state between pDCs and cDCs.

Next, we queried Sincast imputed Villani et al. (2017) data with the reference of the DC and monocyte subset from Rajab et al. (2021) (Supplementary Figure S11). We non-linearly reconstructed the PCA projection landscape with diffusion map, embedding the atlas samples and query cells into new data coordinates of diffusion components (Section 4.7). We found that DC5 cluster projected between conventional DCs and plasmacytoid DCs, suggesting a dual identity (Figure 3C, Supplementary Figure S5). This results was consistent with Villani et al. (2017) who claimed that DC5 represent a new sub-population of DCs which lie on the continuum between these two states. This highlights how Sincast imputation and projection can reveal new cell states which may not exist on the reference data.

#### Sincast imputation can highlight pseudo-time trajectories

We considered a subset of data from Bian et al. (2020) corresponding to macrophages from the embryonic head and york sac. The cells were projected onto Rajab et al. (2021) atlas after Sincast imputation. As expected, the cells were projected close to fetal microglia in the reference (Figure 4A). When we investigated our modified Capybara score for each of the projected cells against the reference microglia cell types, there was an increase of this score according to the Carnegie stage of the embryo (Figure 4B). This result showed that Sincast imputation followed by projection can preserve the inherent time course information in the query data.

**Figure 4.**
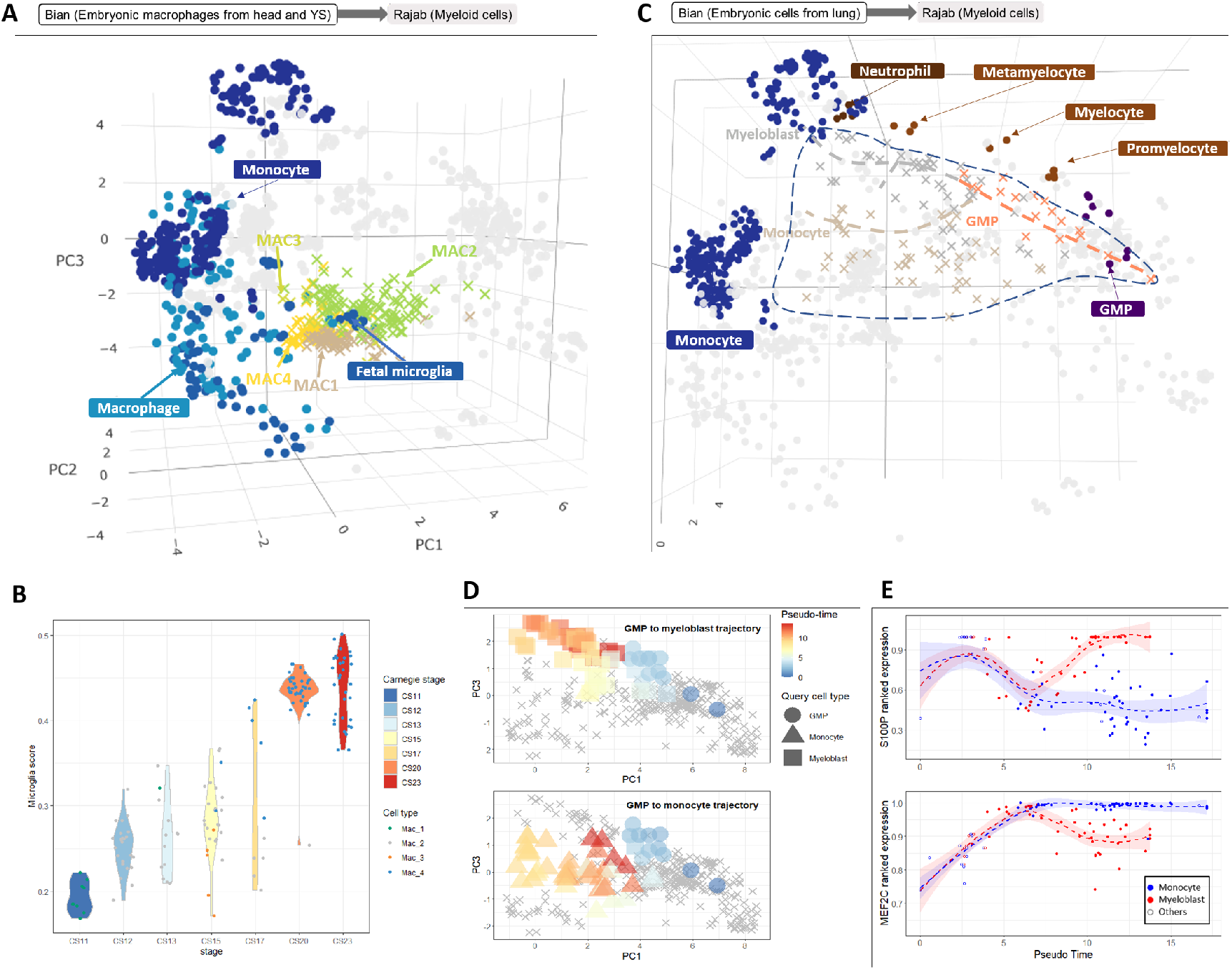
**(A)** Projecting embryonic macrophages from Bian et al. (2020) onto Rajab et al. (2021) after Sincast imputation revealed their identity to be closest to the fetal microglia in the reference. **(B)** Sincast preserved the development trajectory inherent in the query data. Modified Capybara score of these cells against the reference microglia showed increasing values with their Carnegie stages. **(C)** Sincast workflow can produce pseudo-time trajectories. We applied a trajectory inference algorithm (Slingshot) to the projection of another subset of query data from Bian et al. (2020) after Sincast imputation. This showed pseudo-time trajectories from GMPs towards either monocyte or myeloblast fates. **(D)** Projected cells coloured by pseudo-time calculated from (C) showed a clear concordance with the annotated cell types in the query data. **(E)** These trajectories can then be used to find key genes of differentiation. The expressions of neutrophil specific gene S100P (top) and monocyte specific gene MEF2C (bottom) were plotted against the pseudo-time values of the projected cells. These showed clear branching of their expression according to the cell fate.

We then considered a different subset of cells from the same query data, involved in the monocyte to neutrophil differentiation process in the lung, and projected these cells onto the same reference atlas after Sincast imputation. We ran the unsupervised trajectory inference algorithm Slingshot from Street et al. 2018 on the PCA of the projected cells (Figure 4C). This analysis highlighted pseudotime trajectories originating from Granulocyte-monocyte progenitors (GMP) and branching towards the myeloblast and the monocyte cell fates (Figure 4D). When we identified the highly expressed genes for the cells at the end of these trajectories, they represented the typical marker genes that are associated these cell types (i.e. S100P for neutrophil, MEF2C for monocyte) (Figure 4E).

These examples showed that pesudo-time trajectories can be inferred correctly from the Sincast workflow. They also illustrate another major advantage in performing pseudo-time analysis after projecting onto a reference: only subsets of the query data are required, as the reference data already provide sufficient underlying structure for a trajectory analysis. In addition, simpler trajectory algorithms can be used when a bulk reference atlas is used, since complex graphs do not need to be constructed. Another advantage is that formal differential expression analysis is also not necessary to highlight key genes along a trajectory, as these are already specified from the principal components.

## 3 Discussion

The analysis of scRNA-seq data requires unbiased characterization of the transcriptional identity of each cell. Even though many bulk RNA atlases have been developed over the decades – covering most tissue types and offering rich phenotype data such as FACS markers and extensive sample annotations, they have been currently ignored in cell type annotation and cell identity prediction tools. Our computational framework is designed specifically to leverage these well curated and established bulk transcriptional data as references. Sincast projects query scRNAseq data onto the low-dimensional expression space learnt on the bulk reference using PCA. PCA preserves euclidean distances between cells and produces new data coordinates that are easy to interpret, compared to non-linear data embedding methods such as UMAP, and is more suited to bulk data. When projected to the bulk atlas, the transcriptional identity of each single cell can be interpreted visually, based on its location on the atlas, but also quantitatively, using our improved Capybara cell score. Two query data processing pipelines are proposed, aggregation and imputation, to mitigate the structural discrepancy between bulk and scRNA-seq data in the projection result.

Our first approach, cell aggregation, generates in-silico mimics of bulk RNA-seq samples and is primarily designed for recovering pseudo-bulk identities of cell populations in the query scRNA-seq data. Cell aggregation is easy to implement and preserves global scale and genuine population differences of the query data. Moreover, pseudo-bulk samples have valid statistical interpretation as they are built based on bootstrap sampling of query cluster averages. By visualising the degree of overlap between clusters of pseudo-bulk samples on the atlas, one can obtain a first understanding on whether clusters of cells differ significantly based on their averaged expression. Pseudo-bulk analysis is particularly suitable for case-control studies in which cluster level differences are of greater interest than of cellular level variation within clusters, as we showed in Case study 2. An additional use case for pseudo-bulk aggregation is the creation of a reference for evaluation of single cell methods, as we showcased in Case study 4 with the Villani et al. (2017) query for self-projection to evaluate imputation methods. Other use of pseudo-bulk aggregated data include appending an existing bulk atlas to extend its range of cell states. Sincast facilitates this process through its aggregation workflow.

However, aggregation also has its limitations as pooling and averaging ignores within cell cluster variation. As a consequence, meaningful sub-population signal detected by scRNA-seq can be masked in pseudo-bulk samples. For example, our attempt to project the Bian et al. (2020) data was challenged due to the complexity of the study underlying biology (not shown). Continuous time resolution in cell development was lost, and the number of cells with a common combination of biological attributes (cell type, tissue location, development stage) was too small to generate valid pseudo-bulk samples. Therefore, we recommend using the aggregation approach when existing clustering assignment of the data are reliable, and the aim is to benchmark overall cluster identity. Otherwise, our second approach, cell imputation, which can model and retain complex cell-to-cell relationships in the scRNA-seq data is a better alternative.

We compared the performance of Sincast imputation with three other popular scRNA-seq imputation methods: MAGIC, knn-smoothing and SAVER. We imputed the same query data with the methods’ default parameters. The query projections onto the bulk atlases resulted in different data structures and scales depending on how each method models cell-to-cell relationships. This comparison raised the issue that imputation may induce excessive technical artifacts. Thus, choosing a suitable imputation method with appropriate tuning parameters is important, and should be evaluated with the overall aim of the analysis. Sincast imputation has been designed primarily for projecting the single cell imputed data onto bulk reference, but it can also be extended for other types of analyses, such as clustering, differential expression analysis and multimodal atlases.

Query identity profiling was performed using an improved version of the Capybara cell score from Kong et al. (2020), based on restricted linear regression. We chose Capybara for its ability in providing smooth quantitative profiling of single cells whose identity might be between the major cell types and states of the reference. Other tools were considered, such as Machine Learning classifiers, but they tends to assign cells to specific (discrete) reference categories. However, since collinearity between the reference gene expression profiles affects linear regression models, all predicted cell type scores other than the dominant cell type should be considered when characterizing a query cell identity with our prediction tool.

In conclusion, leveraging established bulk transcriptional atlases as reference data for determining cell identity in scRNA-seq data can lead to powerful biological insights. Sincast is an unique toolkit specifically designed for this purpose, and can be used to comprehensively annotate matching cell states as well as discovering new states. Sincast also provides a novel framework for single cell computational method evaluation.

## 4 Methods

### 4.1 Data description

All data were collected from public data repositories, as described in Table 1.

### 4.2 Building a bulk transcriptional reference atlas

We define bulk transcriptional reference atlas as a Principal Component Analysis (PCA) representation of a gene expression dataset to which external data (i.e. scRNA-seq data) can be projected and queried. This section details the data pre-processing steps required to build the reference atlas prior to PCA (Figure 1), where we assume that quality controls on the reference data, such as low quality gene and sample filtering have been performed.

We first perform rank transformation (RT) to normalize the reference data, as previously described by Angel et al. (2020), and further detailed in Supplementary Material 5.1). Only discriminant genes relevant for classifying the reference sample cell types (or any other class of interest) are selected to build the reference atlas (we summarise the number of genes retained in our case studies in Table 1). For data without distinct class assignment, one can either perform sample clustering on the data first, or use highly variable genes as substitute of discriminant genes Yip et al. 2019. We assess the relevance of a gene by calculating the correlation between the samples ranked expression of the gene and the samples (known) cell type labels, using the Hellinger distance (HD). Details on how to calculate the HD score can be found in Supplemental Material 5.2.

Sincast projection requires that the query genes match the set of genes used to construct the PCA reference atlas. Hence, overlapping discriminant genes are retained between reference and query. The reference data is rank then transformed again to adjust for the change of gene sets and the reduction of available ranking allocation. PCA with gene centering is then applied to the reference data to project samples into low dimensional coordinates that maximize sample variation (as detailed in Supplementary Material 5.3).

### 4.3 Projecting the query data onto the bulk reference atlas

We define projection as mapping query cells onto the PCA space of the reference atlas. This allows us to benchmark query biology by measuring the cell locations relative to the distributions of the reference samples from the atlas. Rank transformation followed by gene centering is applied to the filtered query data, where centering factors of the query genes are the same as from those of the reference data. We project the query cells by multiplying their centered rank profiles with atlas gene loading matrix, which defines rotation of gene coordinates to obtain atlas PC basis. Reference samples and query cells can then be visualized jointly on the atlas coordinates, where distances between samples and cells indicate their transcriptional profiles similarity. However, projecting sparse scRNA-seq query data onto bulk atlases is challenging, as RT is not sufficient for sparse data normalisation. The large proportion of tied gene expression and inflated zeros violates the RT assumption of constant gene rankings across batches and libraries. We describe below how Sincast addresses this issue via pseudo-bulk aggregation and imputation on the query single cell data before projection.

### 4.4 Sincast pseudo-bulk aggregation

Cell aggregation has been used in single cell studies to use bulk statistical methods, such as DEseq2 for differential expression testing (Crowell et al., 2020; Love et al., 2014). In Sincast, we recommend using an aggregation approach when the query scRNA-seq data satisfy the following requirements:

1. Cells can be distinctly separated according to clusters. Cellular variation within cell clusters is not of primary interest, and cells are considered as pseudo replicates.
2. The unit of the query data must be additive (e.g. raw UMI count, TPM or CPM transformed data).

For the latter requirement, note that aggregating log transformed counts is equivalent to multiplying counts and then performing log transformation, thus, the resulting aggregated samples do not represent valid bulk identities of cell populations.

We consider query data clustered according to cell types or other combination of identity labels of interest. We denote the number of cells of cell type *t* as *N_t_*. Aggregation is simply performed by sampling cells of cell type *t* with replacement *N_t_* times, and then calculating the average expressions across re-sampled cells on a gene-by-gene basis to create pseudo bulk samples. The sampling bootstrap procedure is repeated *B_t_* times for each cell type *t* independently, where *B_t_* is usually chosen to be at least *N_t_*. Labels of pseudo-bulk samples are inherited from the labels of single cell cluster from which the samples are generated. Bootstrap sampling is often used for inferring sampling distribution of a given statistic. Here, the idea is to infer the sampling distribution of averaged expression profiles of single cell populations.

### 4.5 Existing imputation methods for scRNA-seq

Rank transformation is limited by small library sizes of scRNA-seq, resulting in many tied expressions and zeros to adequately align query scRNA-seq to reference bulk-seq data. One solution to address the structural discrepancy between the query the reference is to impute and smooth values in the query. Here we describe three best performing scRNA-seq imputation approaches (evaluated by Hou et al. (2020)) that were benchmarked in our study. MAGIC (Van Dijk et al., 2018) in particular prompted the methodological development of Sincast.

#### MAGIC

(Markov Affinity based Graph Imputation of Cells)) (Van Dijk et al., 2018) is based on the theory of diffusion map. MAGIC first computes a cell-wise distance matrix for the query data, then converts the distance into a probabilistic similarity measure called ‘affinity’ using adaptive Gaussian kernels. The affinity matrix is row-stochastic normalized into Markov transition matrix, whose entry represents transition probabilities from the row to the column cells. The imputed expression profile of a cell is the weighted average profile of cells within the targeted cell’s neighborhood where the weights correspond to the transition probabilities of the Markov matrix.

The performance of MAGIC can be largely affected by the tuning parameters, primarily the exponent of the Markov matrix, called diffusion time *t*, the cell neighborhood size, knn-max and the bandwidth of diffusion kernel. The affinity between two cells that are not in each other’s knn-max neighborhood, is set to zero, which means that these two cells will not participate in each other imputation. When knn-max is set to a too small value, the imputed scRNA-seq data will retain a high proportion of zero expression value due to small pooling size; When knn-max is set to a too large value – larger than the cell population size, the cell is almost equally imputed by the other cells in its neighborhood, from the same or different types and states. This is a result of a fast decaying rate of the tail of the Gaussian kernel function, where affinities of cells in a pool are small and indistinguishable (Figure 5). The impact of knn-max is further aggravated by increasing the imputation strength using the diffusion time *t* parameter. Our proposed approach described next addresses these limitations.

**Figure 5.**
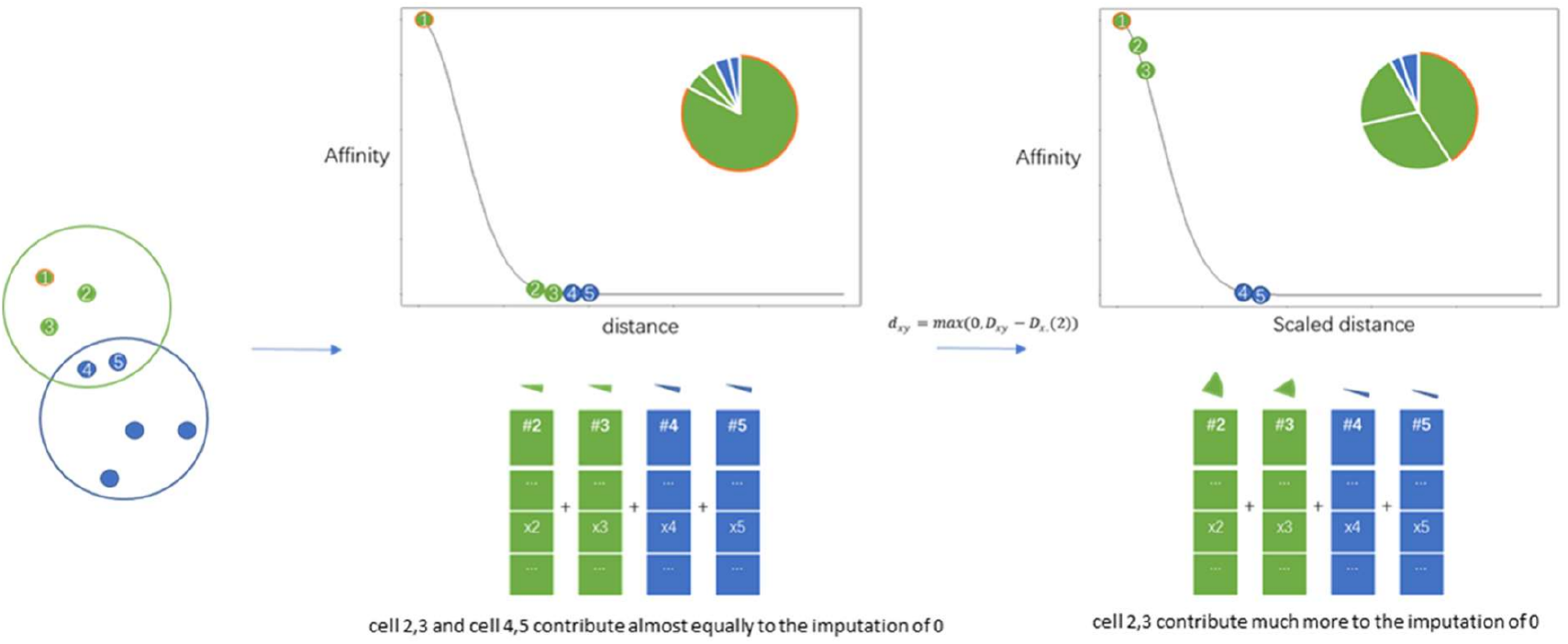
A schematic diagram showing MAGIC sensitivity to tuning parameters. Suppose the query contains two cell populations represented as green and blue points and cell 1 is to be imputed. Using MAGIC affinity matrix specification, cell 4 and 5 contribute highly to the imputation of zeros in cell 1 if a wrong neighborhood size for imputation (5 in this case) is chosen. We propose to address this issue by scaling the distance measurement to create an affinity matrix focuses on cell 1 local connectivity, so that cells 2 and 3 participate more in the imputation.

#### SAVER

(Single-cell Analysis Via Expression Recovery) (Huang et al., 2018) assumes that the UMI counts of scRNA-seq data follow a negative binomial distribution framed as Poisson-Gamma mixture. SAVER performs penalized Poisson Lasso regression of each gene using the rest of the genes as predictors. The fitted regression values are set as prior Gamma means for the Poisson rate, and the Gamma variance is estimated empirically with a maximum likelihood approach. The final imputed value for each gene in each cell is the posterior mean of the Poisson rate, i.e. the weight between the regression fit and the empirical observation.

#### knn-smoothing

(K-nearest neighbor smoothing) (Wagner et al., 2017) first aggregates the expression profile of each cell with its nearest neighbor to initialise the input cells for the next iteration. In the next iteration, the aggregated profiles are smoothed again, but this time each cell is aggregated with its 3-nearest neighbors. The process iterates with increasing aggregation size equals to 2^*i*^ − 1 at *i*^*th*^ iteration. The iteration stops when the aggregation size reaches a set maximum *k*.

### 4.6 Imputation with Sincast: a graph-based approach

Our imputation method is inspired by MAGIC, and is modified on the theoretical basis of diffusion map and UMAP - both are non-linear data embedding methods that recover low-dimensional representation of the manifold underlying data in the euclidean space (Coifman and Lafon, 2006; McInnes et al., 2018; Van Dijk et al., 2018). Our method aims to

1. Infer a *κ*-neighbor graph from the query scRNA-seq data based on UMAP (steps 1-4 in algorithm 1),
2. Construct a diffusion operator from the graph that is applied to the query for data diffusion (steps 5-8 in algorithm 1).

We assume that cells in the query can communicate and exchange their expression profile according to their local arrangement on the manifold. Gene expression of a cell is imputed as the weighted average gene expressions of the cell’s *κ* nearest neighbors. Weight for imputation between a pair of cells is derived from their geodesic distance measured on the manifold. Our pseudo-code is presented in Algorithm 1.

#### Distance scaling

Suppose *G* (Gene) by *N* (Cells) normalised gene expression matrix of the query data *X*. Consider *S* = {*c*_1_, *c*_2_, …, *c*_*N*_} as an ordered set that contains the column vectors of *X*. Cells *c*_*i*_ in *S* are assumed to be sampled from a low-dimensional manifold embedded within the data ℝ^*G*^ expression space. We use a graph 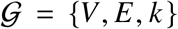 to represent the pairwise geometric relationships of cells on the manifold. In such setting, cells can be considered as nodes of 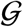 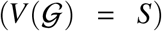, connected by weighted edges, whose weights *W_ij_* are given by the pre-defined kernel functions *k* : *S* × *S* → *R*_⩾0_, *k*(*c*_j_, *c*_*i*_) = *k*(*c _j_, c_i_*). The weight *W_ij_* = *k*(*c*_*i*_, *c*_*j*_) represents the similarity between cells *i* and *j* with respect to their geodesic distance on the manifold, where *k* is derived from adaptive Gaussian kernels applied to pseudo-matrices defined individually for each cell *c*_*i*_ in the query. Denote 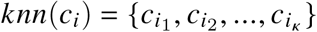 the set of *κ* nearest neighbors of cell *c*_*i*_. As we do not know the true structure of the underlying data manifold, the geodesic distance between *c*_*i*_ and its *j*^*th*^ nearest neighbours *c*_*i*_*j*__ ∈ *knn*(*c*_*i*_) is approximated by the euclidean distance in ℝ^*G*^ (valid only if *κ* is small enough):

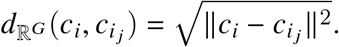

The euclidean distance is then converted to cell-specific pseudo-metrices defined by the distance beyond nearest neighbor:

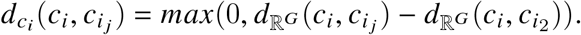

This step of distance scaling can be simplified as follows (for theoretical details, see McInnes et al. (2018))

1. Since now 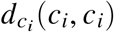 and 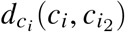 are both 0 and indistinguishable, we can define a graph in which all cells are guaranteed to be locally connected to at least its first nearest neighbor. The weight of self-looping edge (*c_i_, c_i_*) becomes less important compared to the weights of other edges {(*c_i_, c _j_*)| *j* ≠ *i*} connected to *c*_*i*_. As such, neighbors of *c*_*i*_ can contribute more to the inference of *c*_*i*_’s identity, as we illustrated in Figure 5.
2. Because of the curse of dimensionality, the distances between cells in the same neighbourhoor – based on their gene expression, is expected to show little variation relative to the absolute values of distances (i.e. 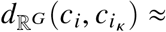 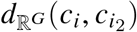. We subtract the distance to each cell’s first nearest neighbor to mitigate that effect in the graph construction, and to put more emphasis on distances differences among neighbors.

#### Weight adjacency matrix

Next, we define the adaptive kernels 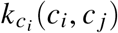 for *c_i_* as follows:

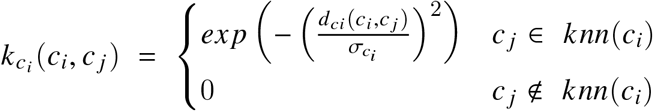

The kernel bandwidth 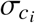 is defined locally for *c*_*i*_ with respect to the *c*_*i*_ cell-specific pseudo-metric such that 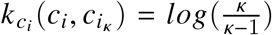. The probabilistic interpretation for the choice of bandwidth is that all the cells in *X* are set to communicate with their *κ*^*th*^ nearest neighbors with a fixed probability equal to 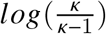. Each cell’s bandwidth is derived from its distance to its *κ*^*th*^ nearest neighbor, which gives a proxy of the cell’s local density. Hence, by normalizing distances with local densities of cells, weights of connection between cells are defined irrespective of sampling density of the data.

We have already obtained a directed graph with asymmetric weighted adjacency matrix *W*^*asy*^ whose entries are given by 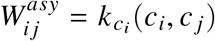. However, asymmetric weights among different cells are not compatible as these weights are computed based on different matrices. To construct a valid Laplacian graph and hence a Markov transition matrix for data diffusion, we define a symmetric *W* based on *W*^*asy*^ to represent the final undirected graph 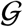:

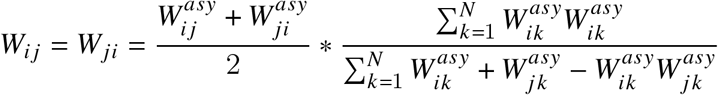

The term on right-hand side of the fraction product represents the Fuzzy Jaccard Index (FUJI, Petković et al. 2020) measured between the knn graphs of cell *i* and *j*. We modified FUJI by swapping the minimum t-norm on the numerator to a product t-norm, and the maximum t-conorm at the denominator to a probabilistic t-conorm. Our graph is constructed to highlight the connection of cells that share common neighborhoods. The connectivity constrain down weights potentially poor connections in the graph, and improve the robustness of the imputation.

#### Data imputation

Using the theory of diffision map, *W* is the diffusion matrix defined by (*S*, *k*). Let 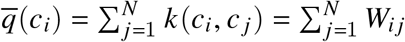 be the finite approximation of kernel volume (or degree in graph) for cell *i*. We define a new kernel scaled by the local volumes for Laplace–Beltrami diffusion,

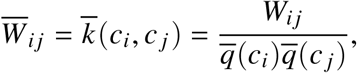

and obtain the Markov transition matrix, or diffusion operator *P* by row stochastic normalization:

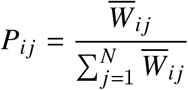

Data imputation is done by applying powered operator *P*^*t*^ on *X*

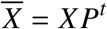

where *t* is a positive scale parameter which controls the step size of diffusion random walk. A large *t* value usually results in stronger imputation strength and less noisy data, but also over-imputation. The risk is a loss of biological signal as the Markov process may attract the identities of minor cell populations towards the regions in 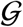 with low escaping probabilities (these regions often correspond to discrete biological niches) in a long-time diffusion.

#### Visualisation

To get a sense of the geometry of the data which defines the graph used for data imputation, we can visualize the data embedding by mapping each cell *c*_*i*_ to its first three diffusion coordinates

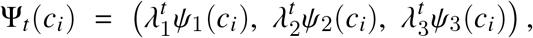

where *ψ*_1_, *ψ*_2_, *ψ*_3_ are the left eigenvectors of ***P*** with the top three largest corresponding eigenvalues 1 > *λ*_1_ ⩾ *λ*_2_ ⩾ *λ*_3_ ⩾ 0. These eigenvalues are only strictly less than 1 if the graph is connected. The constant eigenvector *ψ*_0_ of ***P*** with eigenvalue *λ*_0_ = 1 is not of our interest and so is omitted from the visualization.

#### Parameter tuning

By default, the graph of the query data is computed based the PCA of *X* for dimension reduction and global noise filter prior to distance calculation (see Algorithm 1). By default, *κ* = 30. Two alternative ways of choosing *κ* are also proposed based prior assumption on the characteristics of the data set:

*Option A.* We can approximate the minimum *κ* that gives a connected graph. This approach is recommended when we assume that no cells or biological components in the data are functionally isolated.

*Option B.* We can approximate the minimum *κ* to reduce the sparsity of the data to 25% when *t* = 1. This approach avoids tuning *t* in the imputation, but the euclidean distance may no longer be a valid approximation of geodesic distance when *κ* is large.

In A, if *κ* is much larger than in B, the latter should be preferred.

We found that the parameter *t* had a significant impact on the imputation result, based on our case studies: a large *t* value tended to distort the data structure compared to an imputation with *t* = 1. For most of the query data we examined, a small *t* ≈ 3 was usually enough to reconstruct complex cell-to-cell relationships with a wide range of *κ* values. Regardless of the tuning of our parameters, we showed that our methodological improvements, such as using a distance beyond nearest neighbor and FUJI greatly compensated for a poor parameter choice, highlighting our algorithm’s robustness and accessibility for imputation and method evaluation.

#### Data scaling after imputation

We found that nearest neighbor based graph imputation methods (e.g. MAGIC, knn-smoothing) can easily over-smooth the query data when the tuning parameters are not chosen carefully. For instance, the projection of query data imputed by MAGIC showed strongly reduced local variance and shrinking of the global structure relative to the atlas landscape when the diffusion time *t >* 1. The loss of local variation is expected due to averaging gene expression of cells within each cell’s neighborhood. The shrinkage of query distribution towards its global average happens when the cells’ defined local neighbourhood sizes are larger then their actual size (as we showed when comparing the MAGIC and Sincast in Figure 3). To prevent over-imputation and creating technical artifacts to the query data, we propose a scaling approach to shrink the imputed data back to the original data, and recover part of the lost variance due to imputation. The degree of shrinkage in each cell is determined according to the amount of variation change in data due to imputation.

Briefly, we take the weighted average between each cell’s original and imputed expression profile as the data scaling result. Post-imputation data variance upweights the imputed profile, whereas imputation strength measured by the deviation between the original and the imputed data up-weights the original profile (see Supplemental 5.4 for more details).

### 4.7 Non-linear visualisation projection via diffusion map

After projecting the query data onto reference atlases, we apply diffusion map (DM) (Coifman and Lafon, 2006) to the concatenated PC scores (up to the elbow point) of query cells and reference samples to recover the manifold of the projection landscape. Indeed, we can only and practically visualize the first three PCs fitted on the reference samples, but these PCs only reveal the most important variations related to the reference biology, but not to the query. Query specific but important information beyond the first three PCs can be missed. DM enables a fast, non-linear reconstruction of the projection result, allowing for better visualization.

We used function *diffusion()* from the R package *diffusionMap* (Richards and Cannoodt, 2019). PHATE (Moon et al., 2019), a DM based dimension reduction method can also be an alternative. Diffusion bandwidth in DM is data specific, set to be two times the maximum distance between the reference atlas sample pairs. We chose a large enough bandwidth to avoid creating a disconnected representation of the projection landscape. For a large integrated reference atlas rich in biological heterogeneity, a too small bandwidth only emphasises on the differences between atlas samples with distinct identities, and will make the local views between single cells disproportionally smaller than the global view dominated by the atlas samples. As such, local views of projection will be difficult to visualize.

#### Algorithm 1 Pseudo algorithm for our graph-based imputation method

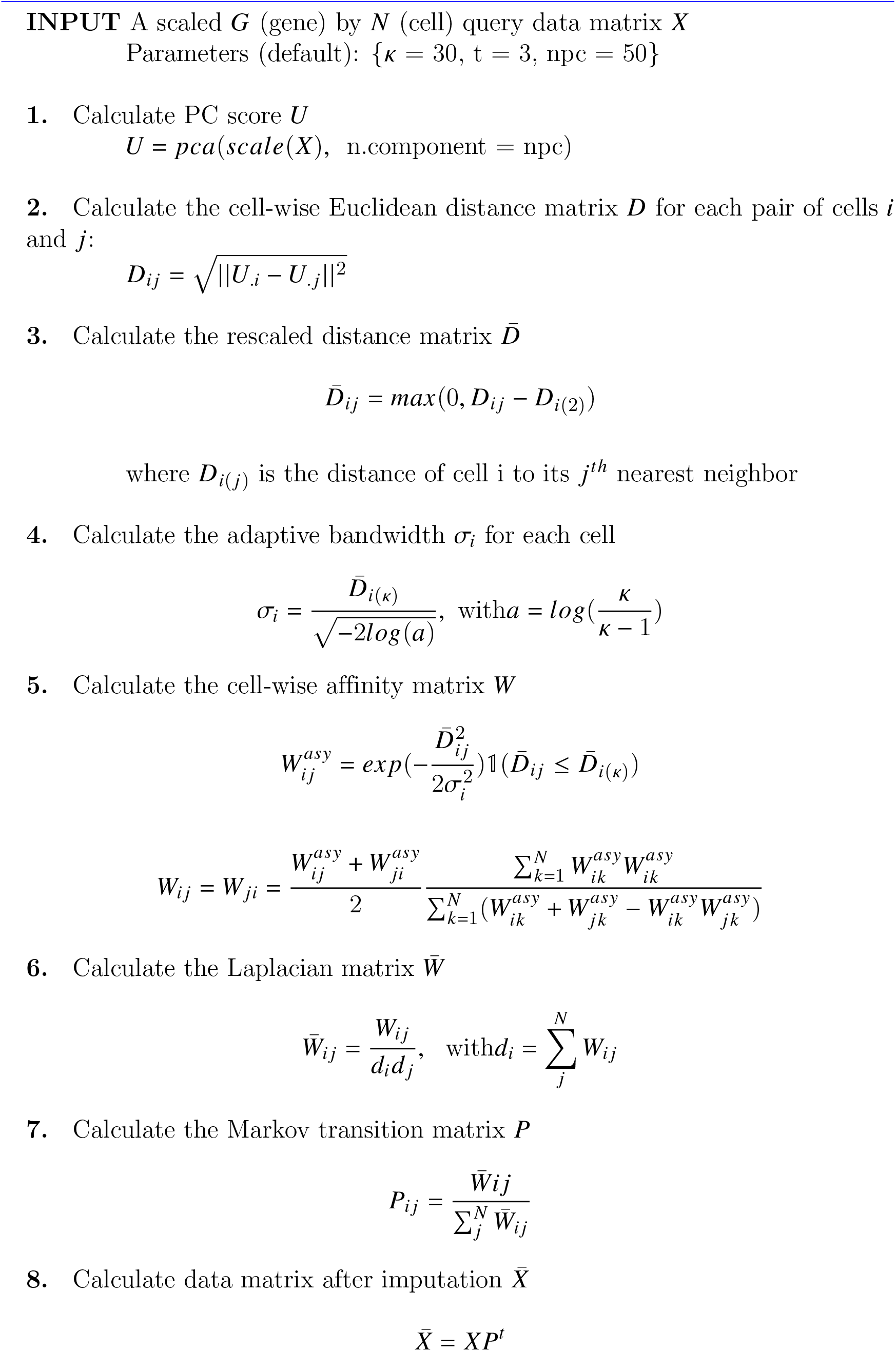

### 4.8 Capybara cell score for continuum cell identity prediction

We applied the Capybara cell score (Capybara, Kong et al. 2020) to predict continuum identities of the query cells. Capybara performs restricted least square (RLS) regressions on each query cell transcriptional profile using cell type, or cluster, averaged expressions of reference samples as predictors. Regression coefficients fitted for each predictor (cell type reference) correspond to identity score predictions. Capybara constraints the coefficient estimates on each query cell to be positive with total sum less than one for biological interpretation. We made two adjustments to improve the predictive performance of Capybara, as described below.

#### weighted RLS

Since different genes may have different degrees of contribution in explaining cell identities, we performed weighted RLS to assign observational weights to each gene corresponding to their importance in classifying cells. These weights (i.e. gene importance) can be estimated from the reference data in a various ways, including standardized gene variance, differential expression p-value, or variable importance metrics from machine learning classifiers. We used gene Hellinger Distances (also used for variable selection to build the atlas).

#### Regression on neighboring samples

To take into account of biological heterogeneity in a comprehensive reference atlas, we propose to regress the query cell expression profiles on their neighboring samples within each atlas clusters, defined as the nearest sub-cluster medoids, rather than on the cluster averages (see Supplemental 5.5 for more details).

### 4.9 Clustering assessment of query observations after projection

We used clustering performance of query projections on atlases as a mean to evaluate the goodness of projection. Clustering performances were quantified using the Silhouette Index, the Distance ratio, and the Adjudted Rand Index (see more details in Supplemental Material 5.7).

## Code availability

Sincast R functions and code can be found on https://github.com/meiosis97/Sincast.

## Acknowledgements

KALC was supported in part by the National Health and Medical Research Council (NHMRC) Career Development fellowship (GNT1159458). We also acknowledge the Discovery Project DP200102903 from the Australian Research Council.

## 5 Supplemental Text

### 5.1 Sample-wise data normalization with rank transformation

We used rank statistics to address technical variation in transcriptomic data analysis, similar to the approaches of Angel et al. 2020; Bolstad et al. 2003; Tang et al. 2021. Here, absolute gene expression values are first transformed to within sample rank percentiles. Previously, we showed that such Rank Transformation (RT) is a simple but robust data normalization technique which can correct for library size differences and account for the presence of technical variation due to sequencing platforms (batch effects) in a dataset that combines independent studies (Angel et al., 2020). RT is applied independently on each sample expression profile. This provides flexibility in appending any extra sample to the reference without re-processing the entire reference dataset. Moreover, RT fits the data of different size and scales, including pre-normalized data publicly available, enabling to customize suitable reference atlases.

RT assumes that technical variation barely changes the relative expression levels of genes within samples (see Angel et al. 2020). We will briefly describe how RT is performed in Sincast. We first rank the absolute expression values of genes within each sample. The gene with the highest expression level in a sample is assigned to a value equals to the total number of genes in the data, denoted *G*. The gene with the lowest expression level is assigned 1. Ties in expression were equally ranked as if they were the lowest member of the tie. All rank values *R*_*ij*_ for gene *i* and sample *j* are then scaled to (*R*_*ij*_ − 1)/(*G* − 1) so that gene expression values are distributed across [0 1] in each sample.

### 5.2 HD score to identify discriminant genes

We first discretize the gene ranks into *T* categories, where *T* denotes the number of unique cell types and is defined using gene-wise k-mean clustering. This facilitates comparisons between gene expression and cell type distributions using metrics developed for categorical attributes. One such metric is the Hellinger distance (HD), a measure of divergence between two probability distributions (Cieslak et al., 2012; Fu et al., 2020). Cieslak et al. (2012) proposed to use HD in binary decision tree to determine the optimal tree split. A good split can create child nodes on which two class labels separate distinctly with little affinity shares between distributions whereby nodes are considered as the support. HD is calculated between the distributions of class labels: the higher the HD, the purer the nodes, and the better the split. In Sincast we use HD to quantify the purity of sample cell types on the unsupervised partitions of genes (analogue to nodes splitting) to assess the genes’ predictive ability.

We calculate the genes HD scores for each cell type using one (labelled class +) versus the rest (labelled class) approach. Consequently, each gene is assigned *T* HD scores that represent its ability in predicting *T* cell types. A gene’s relevance in classifying cell types is defined its mean HD score. Formally, for each gene *g* HD score for cell type *t* is:

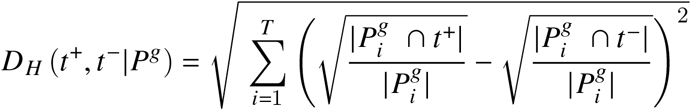

where *t*^+^ is the set of samples labelled with cell type *t*, and *t*^−^ denotes the rest of the samples, 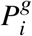 is the *i*^*th*^ partition of gene *g*. By default we select the top 2000 ranked genes with the highest mean HD scores as markers to build atlases. Mean HD scores are also used as gene weights for the improved Capybara cell identity prediction, as described in Section 4.8.

Other metrics such as Information Gain or Gini Index could have been chosen, but are sensitive to class size imbalance and may result in gene selection bias.

### 5.3 Building a bulk reference atlas with PCA

PCA with gene centering is applied on the processed reference data. PCA generates a loading matrix that specifies the rotation of gene coordinates to define the atlas spanned by the principal components (PC) basis. The PC basis represents the latent dimensions embedded in the gene expression space that can capture maximal variations of the samples. Locations of samples in those dimensions known as component scores were computed by multiplying the loading matrix with the data matrix. As a result of matrix multiplication, linear combination of genes realizes rotation and creates components. The number of latent components to include in the atlas is automatically determined by the elbow method, which consists of finding the points of maximum curvature (elbow point) of the changes in cumulative explained variance of PCA with increasing dimensions. The ‘elbow point’ indicates that adding extra components does not significantly increase the variance explained in the data. However, to prevent missing potential subtle biological variations, five components after the elbow point were also added in the atlas.

### 5.4 Post imputation data scaling

First, for a given gene *g* and cell *i* in the imputed data *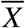*, we scale its expression value with gene-wise scaling factors *f*_*g*_:

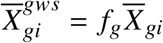

where

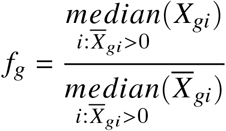

The scaling factors are chosen so that for each gene, cells with observed expressions can roughly retain the same location (in the statistical sense) after imputation. This first scaling step is only valid when both *X* and 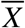 are non-negative, and zeros in the data represent missing values of expression.

Second, we assume that the uncertainty (or variability) of 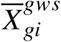 is regularized by the gene *g* underlying mean of imputed expressions. The form of the regularization is modelled as a global dispersion trend of 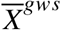, inferred by fitting a generalized additive model (GAM) on the log-transformed gene-wise mean 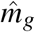, and variance 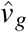 estimations. We assume that 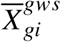 is generated from a gene-wise truncated normal distribution bounded below, where zeros in the imputed data are modeled as weak biological signals that cannot be captured by sequencers and hence censored from observation. The estimation of 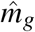 and 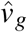 is calculated based on quantile-quantile regression minimizing the following objective function:

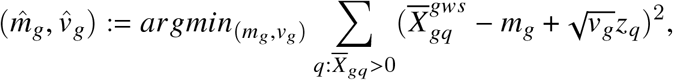

where 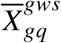 and *z*_*q*_ are *q*^*th*^ quantile for 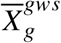 and standard normal *Z* respectively. We use the *mgcv* R package (Wood, 2011). Our GAM model has the following specification:

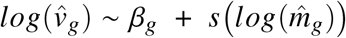

where *s* is the smooth function that defines the cubic regression splines. To prevent over-fitting, the basis dimension of the regression spline is chosen as the smallest value which passes the test of *k.check()* function in *mgcv*. Regression weights 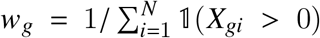 are also considered in this model to account for the uncertainties in model estimation induced by the sparsity of the data. Genes with high sparsity will participate less in the estimation of the global dispersion trend (See Supplementary Figure S14 for trends estimated).

Finally, for 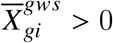, the observational variance 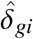 is then approximated by the fitted global dispersion trend as

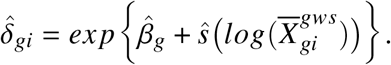

We define the squared imputation residual in each observation as

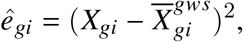

which represents the strength of imputation on the observation. The weighted average between *X*_*gi*_ and 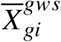 is the final output, and is defined as

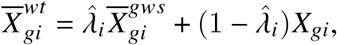

with

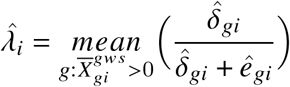

The weight 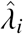 is the mean estimation of imputation’s impact on gene variation in cell *i*, and can be interpreted as follows: When 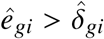, the weighting procedure favors the original data due to a large imputation strength and small post-imputation data variance, suggesting over-smoothing. When 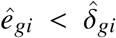, which is typically the case for genes with high sparsity, scaling encourages the imputation of zeros. Supplementary Figure S15B shows that our Sincast data scaling also works on MAGIC imputed data. 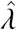 also provides a good indication regarding potential over-smoothing (Supplementary Figure S15C,D).

### 5.5 Capybara cell score

For each cell type cluster *t* of size *n_t_* in the atlas, we first perform Partition Around Medoids clustering on the atlas PC basis to find 5 partitions of the cluster represented by the partition medoids. Each sub-cluster, therefore, contained *n_t_/*5 cells in average. Given a query cell *c* projected onto the atlas, we then searched for its nearest medoid in *t*. Back to the high dimensional space, we denote the ranking expression of the nearest medoid found for c as 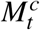. A cell specific reference matrix 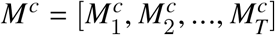 was built, where *T* represents the number of reference cell type to benchmark the query against. Let *y*^*c*^ be the ranking expression of *c*, our goal was to solve the following least square problem,

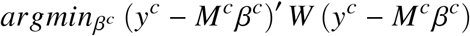

subject to

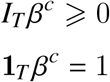

where ***I***_***T***_ is a identity matrix of size ***T***, **1**_*T*_ is a row vector of ones, and *W* is a diagonal weight matrix of size *G*, whose diagonal entries incorporate gene importance to the estimation of *β*^*c*^. Denote the objective function above as *f* (*β*^*c*^), we observed that

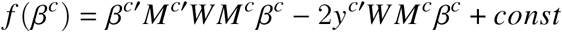

so that

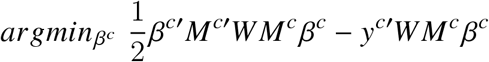

subject to the same regression constrains is equivalent to the least square problem. The updated objective function is in the form of quadratic programming, and so can be solved explicitly. The solution 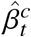 gave the Capybara cell score for cell *c* on cell type *t*.

### 5.6 Down sampling bulk samples to simulate sparse rank expression

We assume that when the total genetic material in a bulk sample reduces, the observed absolute expressions of genes in a sample gradually decreases to zero. Hence we generated the rank profile of a sample with a sparsity rate of *p/G* by substituting the ranks of the first *p* lowest expressed genes to 0, while keeping the rest of genes ranks unchanged. The sparse samples generated were then interpreted as pseudo-single cells. However, our simulation procedure cannot model variation loss in single cells due to rank ties that were not zero expressions. To quantify such, we propose an *effective sparsity* metric, as we describe next in Section 5.7.

### 5.7 Metrics

#### Effective sparsity

Let *R*_*ij*_ be the *i*^*th*^ rank of *i*^*th*^ gene in cell *j*. The effective sparsity of cell *j* is defined as

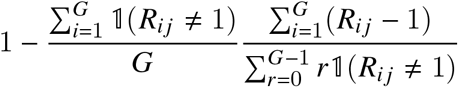

where the left of the fraction product calculates the proportion of gene expressed in cell *i*, adjusted by the factor on the right whose numerator calculates the observed sum of ranks in cell *j*, and the denominator gives the maximum possible sum of ranks on expressed genes when all of them are distinctly expressed.

#### Silhouette index

*Silhouette index* (SI) measures how well cells were separated according to a known clustering assignment (Rousseeuw, 1987). Clustering assignments of query cells were assigned the original cell type labels from the query study.

For each projected cell *i*, we measured its average inter cluster distance

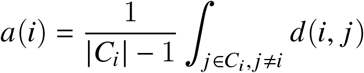

where |*c*_*i*_| is the size of the cluster to which *i* belongs, and *d*(*i, j*) is the Euclidean distance between cells *i* and *j* from the same cluster calculated on the atlas’ PCs.

We also measured the minimum average intra cluster distance of cell *i* as:

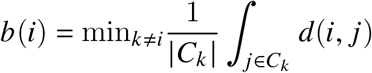

where *C*_*k*_ is a cluster which is distinct from *C*_*i*_. The SI for cell *i* is then defined as

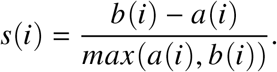

SI ranges between [-1, 1] with larger value indicating that a cell is more similar to the other cells within the same cluster than the cells outside that cluster. In Section 2.2 for Case study 4, SI for each query cell was calculated with respect to the atlas samples because the query true atlas correspondence was known for the pseudo-bulk atlas. The *i* index represents the target query cell, {*j*|*j* ∈ *C*_*i*_, *j* ≠ *i* are the samples from the same cluster as *i* in the atlas and {*C*_*k*_|*k* ≠ *i*} represent the remaining atlas clusters.

#### Distance ratio

When assessing the impact of sparsity on query projection in Section 2.1, we defined the minimum inter cluster distance as

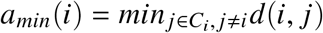

and the minimum intra cluster distance as

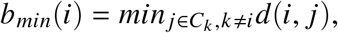

using the same notations as in the SI calculation. Distance ratios of pseudo-single cell *i* were then calculated by

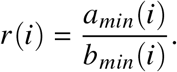

We chose the minimum distance as the measure of cluster affinity to account for heterogeneous within-cluster variation in bulk atlases.

#### Adjusted Rand Index

*Adjusted Rand Index* (ARI) was used to calculate the concordance between two sets of cluster assignments (Hubert and Arabie, 1985). We performed cluster analysis on top PCs of the projected query cells and compare the result with the query’s original cluster labels obtained by the query study. We chose k-mean clustering analysis, where k was set to be the same as the number of original query clusters. Rand index (RI) was then calculated between the two sets of cluster assignments, and was adjusted to correct for the possibility of obtaining the RI by chance. ARI ranges from 0 to 1 with a larger value suggesting that the k-mean clustering of projected observations resembles the query’s original cluster assignments. Hence, we recruited ARI to quantify whether unsupervised clustering methods represented by k-mean clustering can still perform well on the projection of imputed query data. K-mean clustering was implemented using the ‘eclust’ function from the R package `factoextra` (Kassambara and Mundt, 2017). ARI was calculated using the ‘cluster.stat’ function from the R package `fpc` (Hennig, 2020).

### 5.8 Querying Jurkat cell line on the Schmiedel atlas

To test Sincast when the query and the reference data are not compatible in biology, we build an atlas with the reference data from Schmiedel et al. (2018), and we queried the pseudo-bulk sample aggregated by single cells of the Jurkat-T cell line from 10x Genomics (Zheng et al., 2017). We also used two additional bulk RNA-seq Jurkat samples (from FANTOM5 (Lizio et al., 2015) and ENCODE projects (Davis et al., 2018), identity: ENCSR000BX) as query data acting as further benchmarks for the pseudo-bulk projection (see also Table 1).

Without batch correction, query Jurkat single cells, bulk and pseudo-bulk samples were projected to the same region at the middle of the Schmiedel et al. (2018) atlas, being away from all the atlas clusters (Supplementary Figure S1A). Such query arrangement suggests that the query data shares limited transcriptional variation in common with the reference data. We used Capybara cell score to quantify the query identity (Supplementary Figure S1B). In average, the reference could only explain about 25 percent of variation in raw, unaggregated single cells. In contrast, the pseudo-bulk sample had about 60 percent of variation explained, same as which of the real bulk samples. In either cases of raw single cells, bulk and pseudo-bulk samples, the majority of the query variation was explained by the Activated naive CD8+ T cells (CD8-*T A*) of the reference. Enrichment analysis showed that CD8-*T A* was enriched in biological process and functions related to chemokine receptors, cytokine production, and T cell signaling (Supplementary Figure S1D). Characteristic genes of Jurkat T cells such as Interleukin 2 (IL2) were in the top list of differentially expressed genes of CD8-*T A* (Supplementary Figure S1C). Thus, CD8-*T A* is a good reference vocabulary representing Jurkat T cells. Consistency of profiling results on bulk and pseudo-bulk samples suggests that Sincast prediction is robust to query data’s batch sources. When the query and the reference data are not consistent, Sincast can pick out the best reference vocabularies characterizing the query while quantifying the prediction uncertainty.

### 5.9 Impact of sparsity on query projections

The maximum allowed sparsity rate (MAS) for scRNA-seq query projections is atlas and cell type specific. We demonstrate that the MAS for querying on an atlas can be approximated empirically by down-sampling the reference samples’ expression profiles, simulating sparse samples that are then projected and the projections compared with their original locations on the atlas. Down-sampling was performed by gradually substituting ranks of genes in a sample to zero by the order of gene rank (Supplementary Material 5.9). We first analysed the Monaco et al. (2019) atlas built for the Ren et al. (2021) query task as an example (Section 2.1, Case study 2). Sparse samples were generated at sparsity rates from 0 to 100 percent with a step of 5. We treated sparse samples for each sparsity rate as pseudo-single cell queries with known atlas correspondence. Centroids of pseudo-single cell cell type clusters were depicted on the atlas in Supplementary Figure S6A. The trajectories of the centroids with various sparsity rate indicated identity shifts of down-sampled bulk samples. Separations of clusters were enhanced when sparsity rate was less than 40 percent. As sparsity rate increased, clusters tended to gather towards the centre of the atlas, indicating a loss of cell identity signal. We assessed the cluster trajectories quantitatively with each pseudo-single cell’s minimum inter-cluster distance to minimum intra-cluster distance ratio, where the cluster of a pseudo-single cell is defined as its corresponding atlas cluster (Supplementary Material 5.7, Supplementary Figure S6B). Cluster trajectories were welly resembled by the distance ratio, which increase, decrease and then increase on the heatmap with increasing sparsity, describing how clusters moved away, back, and then away respectively. For the Monaco et al. (2019) atlas, the suggested MAS where 90 percent of the sparse sample retain a distance ratio greater than 0.5 was 15 percent. For the other atlases, the MAS varied between 10 and 15 percent (See Supplementary Figure S7, S8 and S9).

Here we only assessed the impact of sparsity on single cell projection. Small gene counts are also enriched in single cells, and may technically affect the projection results due to their ties in gene ranking. The observed sparsity of a cell should therefore be adjusted by the cell additional variation loss in gene expression rank ties. We proposed effective sparsity to account for such variation loss (Supplemmentary material 5.7, Effective sparsity). As an example, the raw single cell data of Ren et al. (2021) were projected accurately on the simulated cluster trajectory of the Monaco et al. (2019) atlas (Supplementary Figure S6A). Effective sparsities of the query cells matched the sparsities of the simulated clusters to which the cells were projected, suggesting that the deviation of the query projection from the atlas clusters was driven by sparsity, not biology.

### 5.10 Sincast and MAGIC sensitivity to parameter tuning

As described in Supplementary Material 4.5, knn-max has a large impact on MAGIC imputation performance. We illustrated this issue with Villani et al. (2017) data by sub-sampling the data to 10 DC6 (phenotypically pDC) and 285 Mono1/Mono2 subset (phenotypically classical/non-classical monocytes) (Figure 3.A). We fixed knn-max to 15, and performed MAGIC imputation with a grid of *t* increasing from 1 to 50 with a step of 1. DC6 imputed at each *t* were projected onto the Rajab et al. (2021) Mono-DC sub-atlas, and the centroid of each projection were shown. The result demonstrates that the identity of DC6 population was clearly distorted after MAGIC imputation. As *t* increasds, the DC6 cluster centroid was projected away from the atlas pDC niches towards the monocyte niches. However, after increasing the DC6 population to 20, the imputed DC6 cells were projected closer to the atlas pDC niches, suggesting that the DC6 population identity was recovered (Figure S13).

To mitigate the impact of poor tuning on imputation, we modified the knn-graph construction in MAGIC based on the theory of UMAP (Figure 3). We also proposed post-imputation data scaling to shrink the imputation result back to the original observations and prevent over-smoothing (see section 4.6 for a detailed description). We performed similar analyses as described above for Sincast method with local neighborhood size set to 15 (*κ* = 15, equivalent to knn-max = 15 in MAGIC). We observed that the shifts of the DC6 clusters towards the atlas monocyte population were greatly restrained even when the parameter tuning was mispecified (Figure 3.A).

## 6 Supplemental Figures

**Figure S1.**
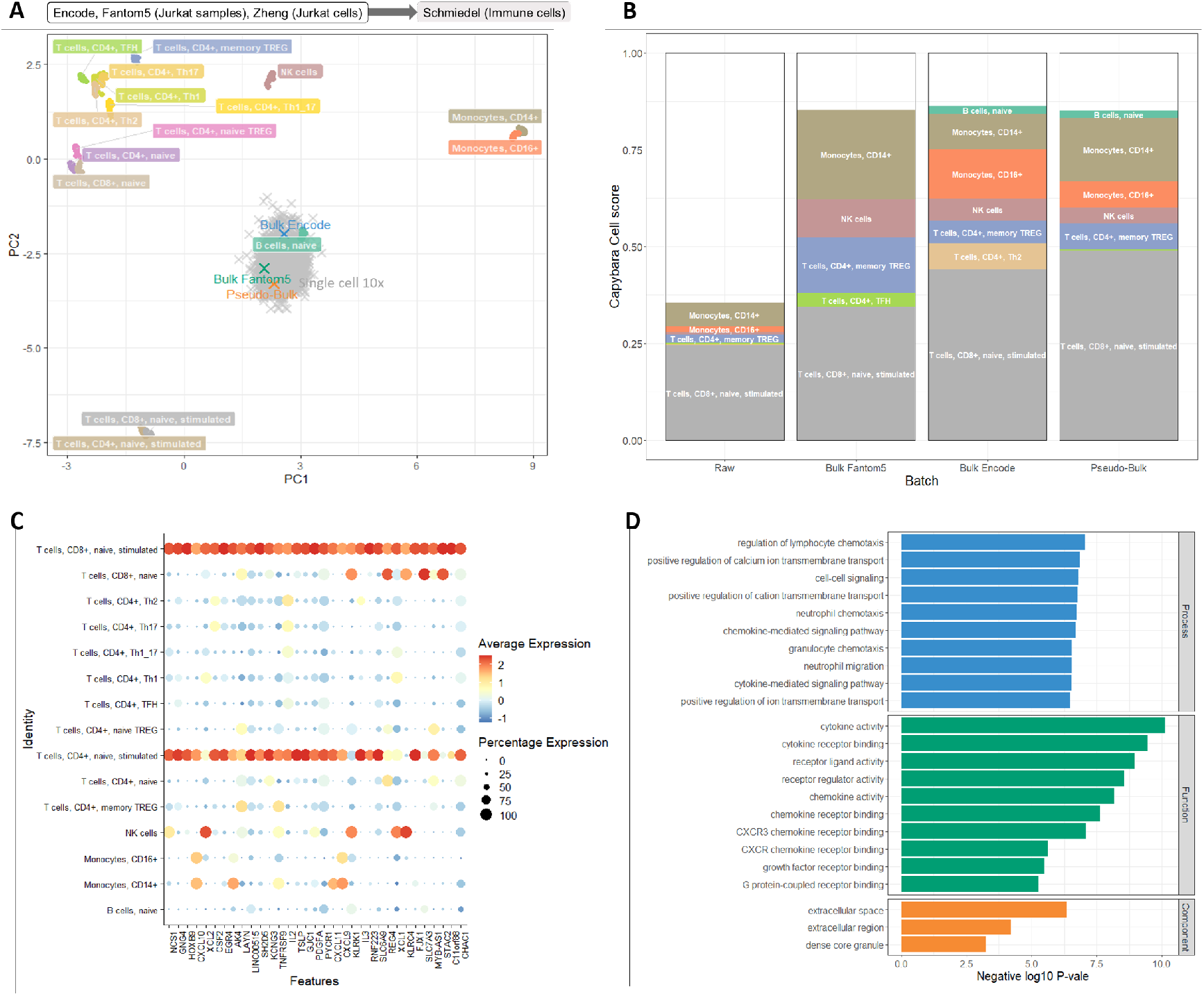
Profiling Jurkat T cell identity on the Schmiedel et al. (2018) atlas. **(A)** On the atlas PC1 and PC3, projection of Jurkat single cell line from Zheng et al. (2017), Bulk Jurkat samples from Fantom5 project (Lizio et al., 2015) and Encode project (Davis et al., 2018), and pseudo-bulk samples aggregated by Jurkat single cells. **(B)** Averaged capybara cell score predicted on the query projections in (A). Heights of the color bars represent the scores query obtained on the different reference cell types. Pseudo-bulk and real bulk samples of Jurkat cell line shared similar transcriptional identity as revealed by their cell score composition. **(C)** We performed differential expression test on the stimulated naive CD8 T cells (CD8-TA) of the atlases versus other atlas samples. Expression of top 36 differentially expressed genes in the atlas samples shown as dotplot. CD8-TA marker gene set overlaps with known markers of Jurkat T cells. **(D)** Gene Ontology enrichment analysis performed on genes from (C).

**Figure S2.**
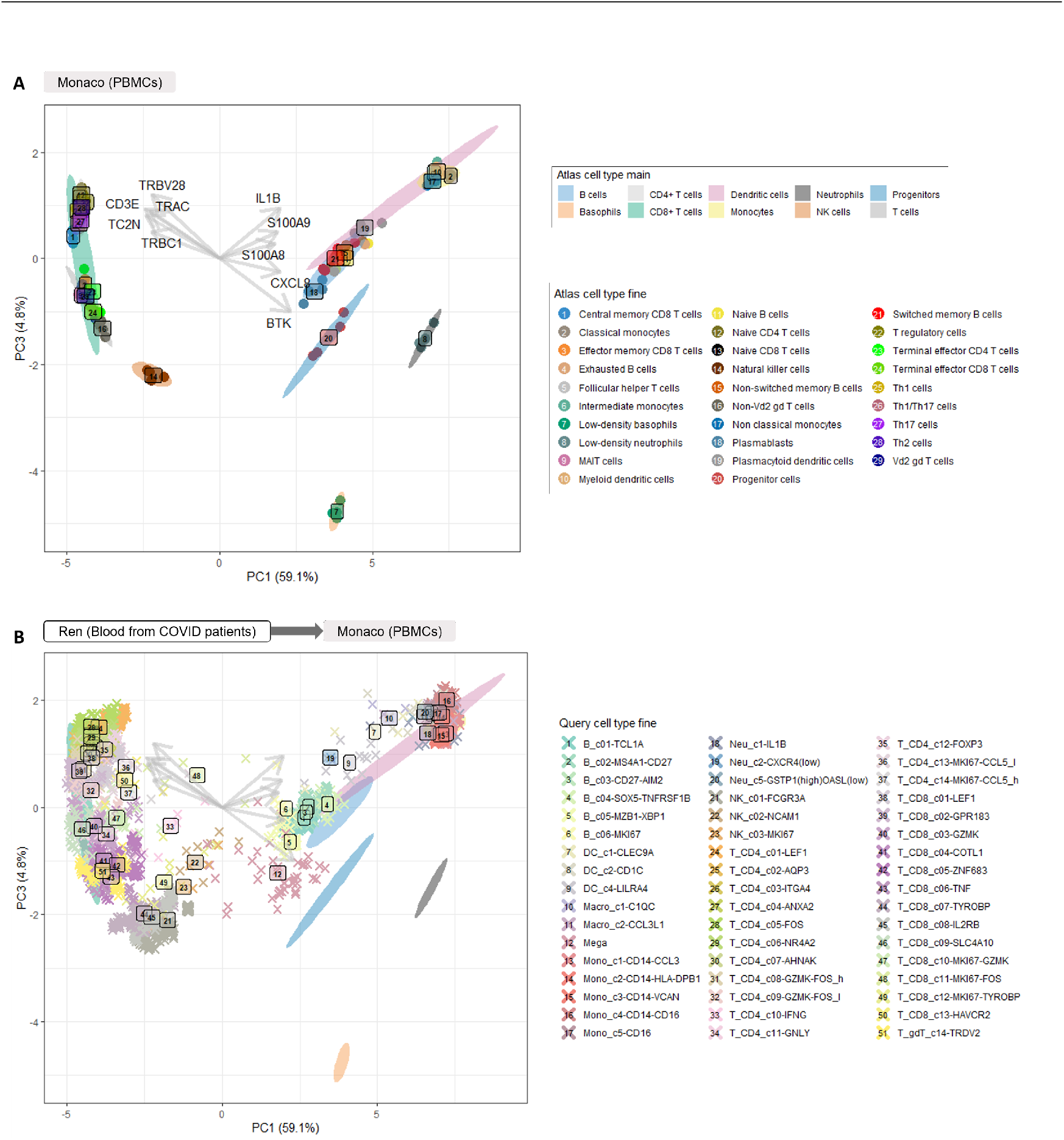
Querying Ren et al. (2021) on the Monaco et al. (2019) atlas. **(A)** Samples from the reference atlas, annotated by the atlas fine cell type labels. **(B)** Query pseudobulk projected on the reference atlas, and annotated by the query fine cell type labels.

**Figure S3.**
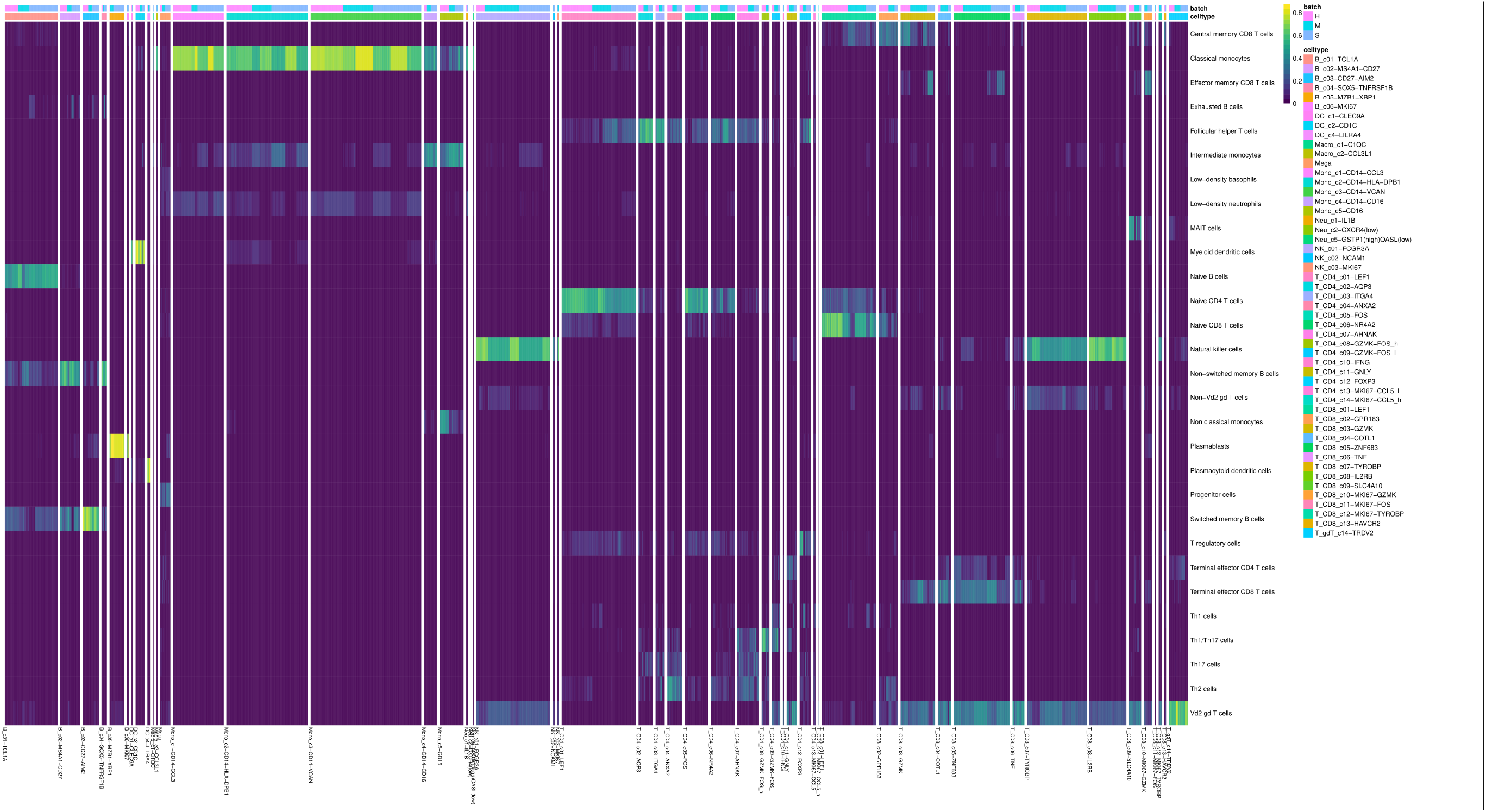
Improved Capybara cell score heatmap predicted on pseudo-bulk samples from Ren et al. (2021) projected onto the reference Monaco et al. (2019) atlas.

**Figure S4.**
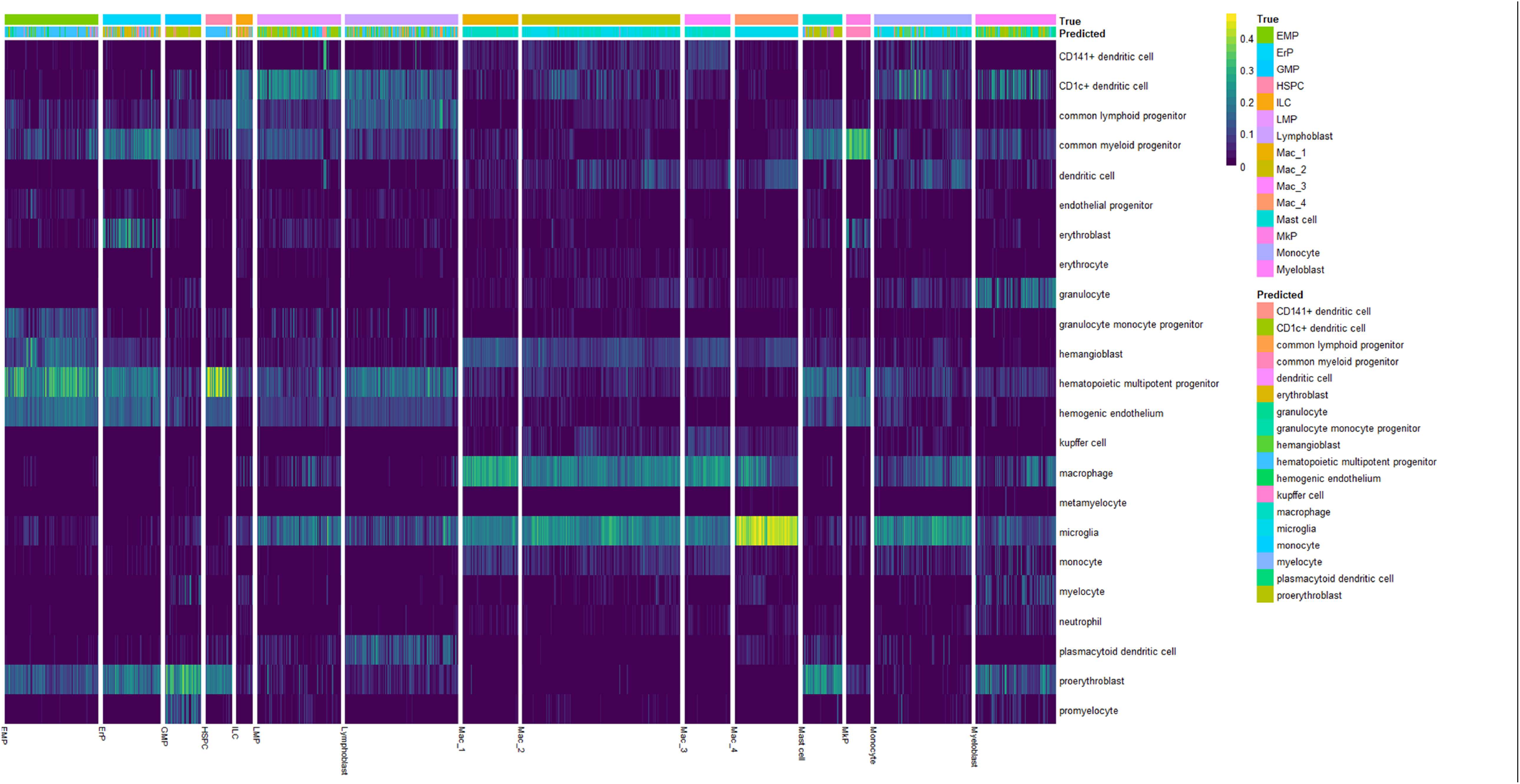
Improved Capybara cell score heatmap predicted on Sincast imputed cells from Bian et al. (2020) projected onto the reference Rajab et al. (2021) atlas.

**Figure S5.**
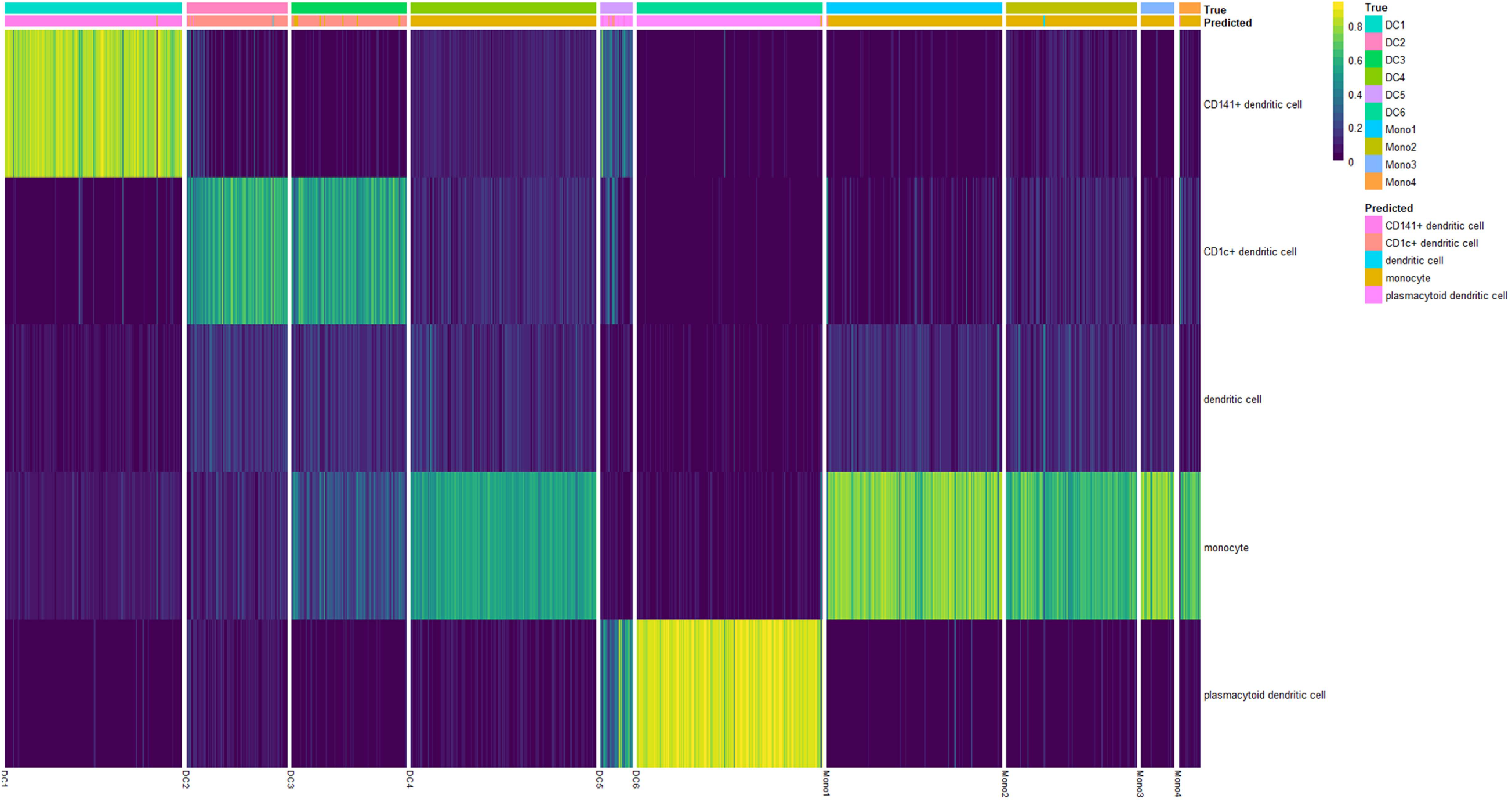
Improved Capybara cell score heatmap predicted on Sincast imputed cells from Villani et al. (2017) projected onto the reference Monocyte-Dendritic cell subset of the Rajab et al. (2021) atlas.

**Figure S6.**
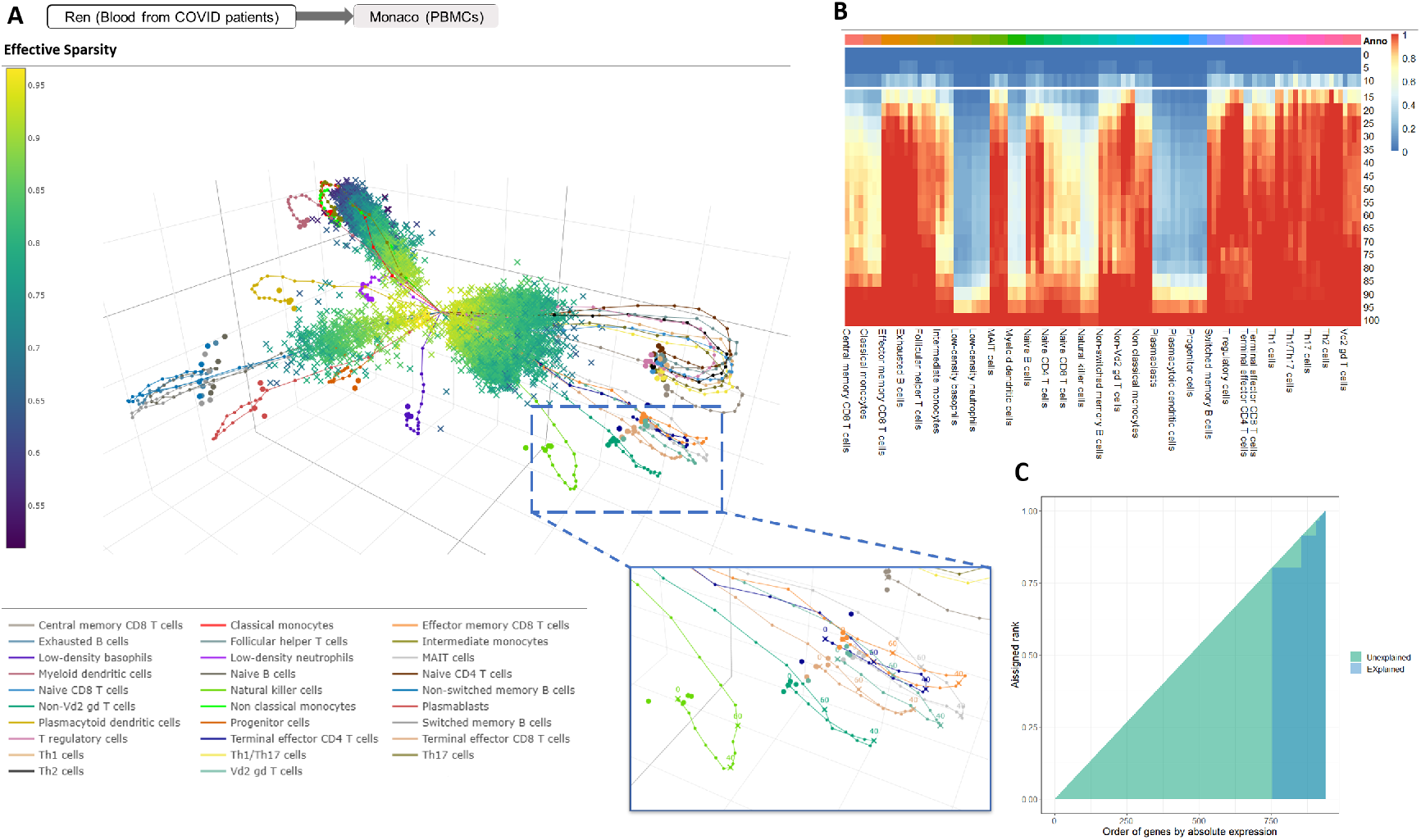
Impact of sparsity on projection. **(A)** Non aggregated query cells from Ren et al. (2021) projected on the Monaco et al. (2019) atlas, and colored by effective sparsity rate. We down-sampled bulk samples in the atlas for sparsity rates ranging from 0 to 100 percent with a step of 5 to simulate pseudo-single cells. Projection of these pseudo-single cells for each sparsity rate are shown as cluster centroids trajectories’. The sparsity of query single cells matched the sparsity of the pseudo-single cell clusters around which query cells were projected, suggesting that deviation of query clusters from the atlas clusters is due to sparsity, rather than biology. **(B)** Minimum inter-cluster distance to minimum intra-cluster distance ratio calculated on the Monaco et al. (2019) pseudo-single cells simulated for each at each sparsity rate. The cluster of a pseudo-single cell is defined by its corresponding atlas cluster whereby down-sampling has not been performed. The change of distance ratio for each cluster matches the simulated cluster trajectories. When sparsity is less than 15 percent, 90 percent of pseudo-single cells retained a distance ratio smaller than 0.5. Hence, querying single cells on the Monaco et al. (2019) atlas may not be largely impacted when sparsity of the query data is less than 15 percent. **(C)** Area plot showing the rank expression of genes within a query cell. On the x-axis, genes are ordered by their absolute expression rank. The blue region depicts the observed ranking. If all genes are distinctly expressed with no ties, the blue region should fill the whole triangle. The green region represents loss of variation in the cell due to ties in expression. Gene rank expression in a cell is not only influenced by sparsity, but also ties in gene expression.

**Figure S7.**
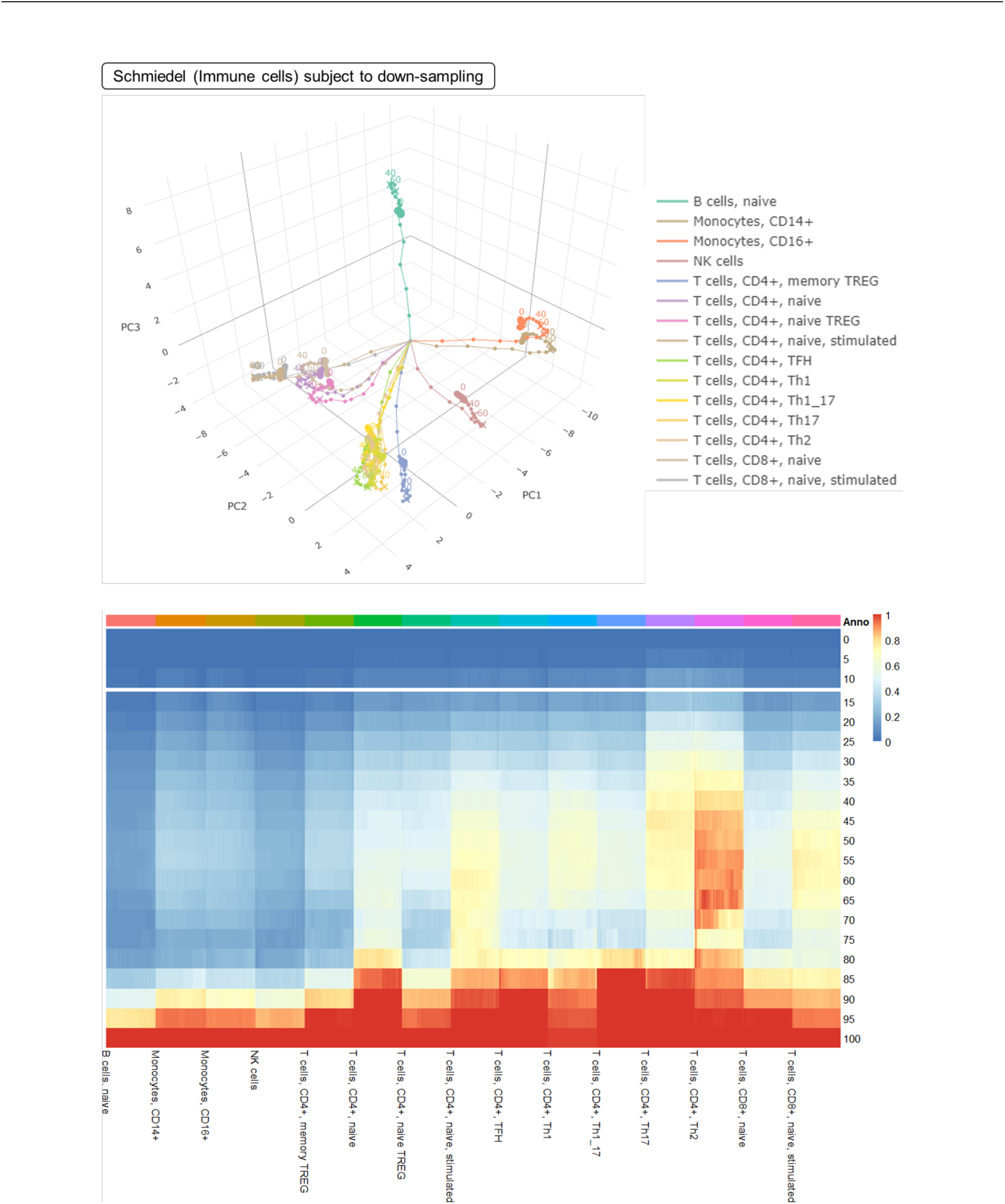
Impact of sparsity on projecting cells to the Schmiedel et al. (2018) reference atlas. Similar to Figure S6 we down-sampled bulk samples in the atlas at sparsity rates ranging from 0 to 100 percent. The top panel shows the atlas and its simulated cluster trajectories. The bottom panel shows each simulated cell’s minimum inter-cluster distance to minimum intra-cluster distance ratio. When sparsity is less than 15 percent, 90 percent of pseudo-single cells retained a distance ratio smaller than 0.5. Hence, querying single cells on the Schmiedel et al. (2018) atlas may not be impacted when the sparsity of the query data is less than 15 percent.

**Figure S8.**
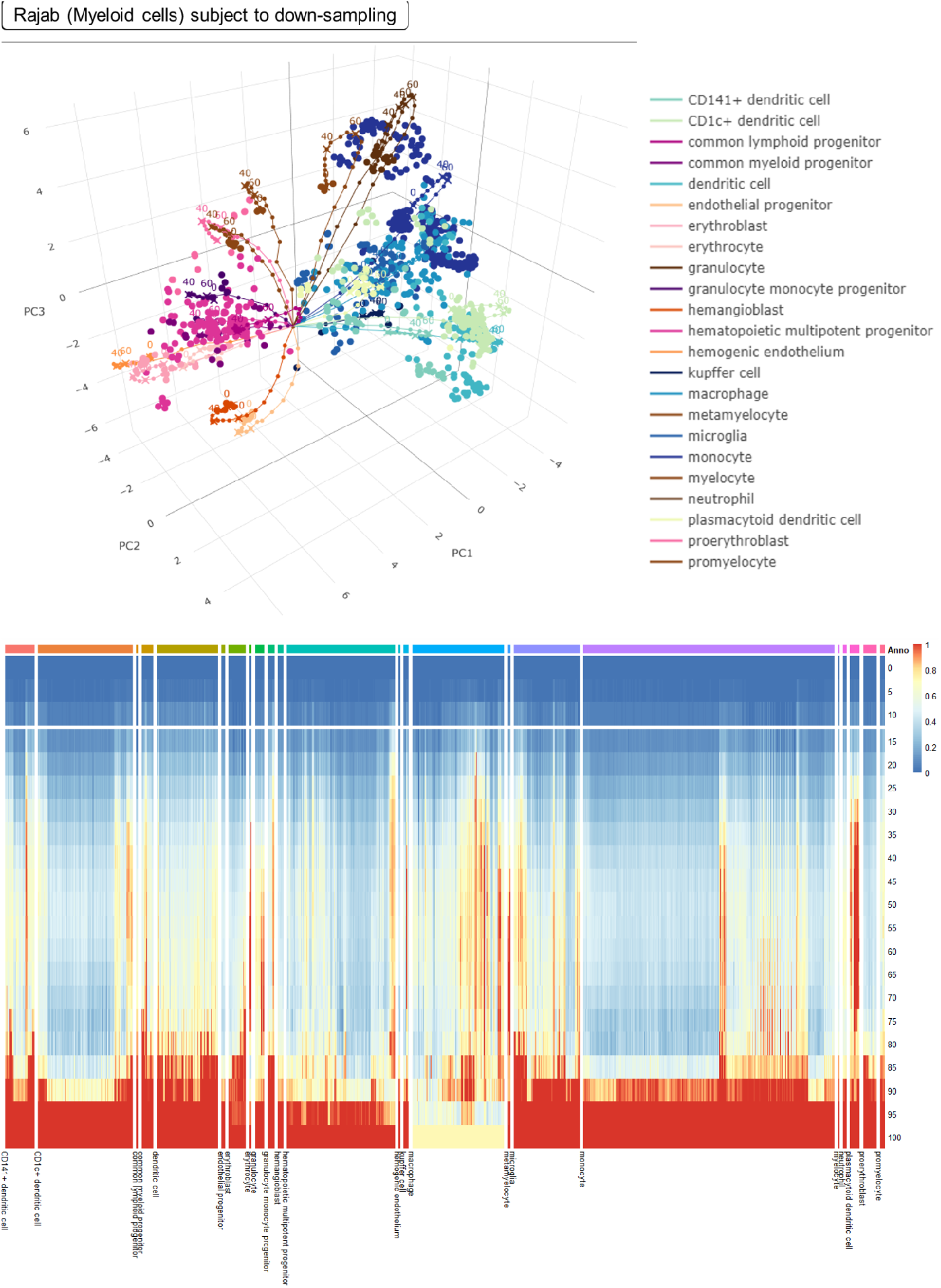
Impact of sparsity on projecting cells to the Rajab et al. (2021) reference atlas. Similar to Figures S6 and S7 we conclude that querying single cells on the Rajab et al. (2021) atlas may not be impacted when the sparsity of the query data is less than 15 percent.

**Figure S9.**
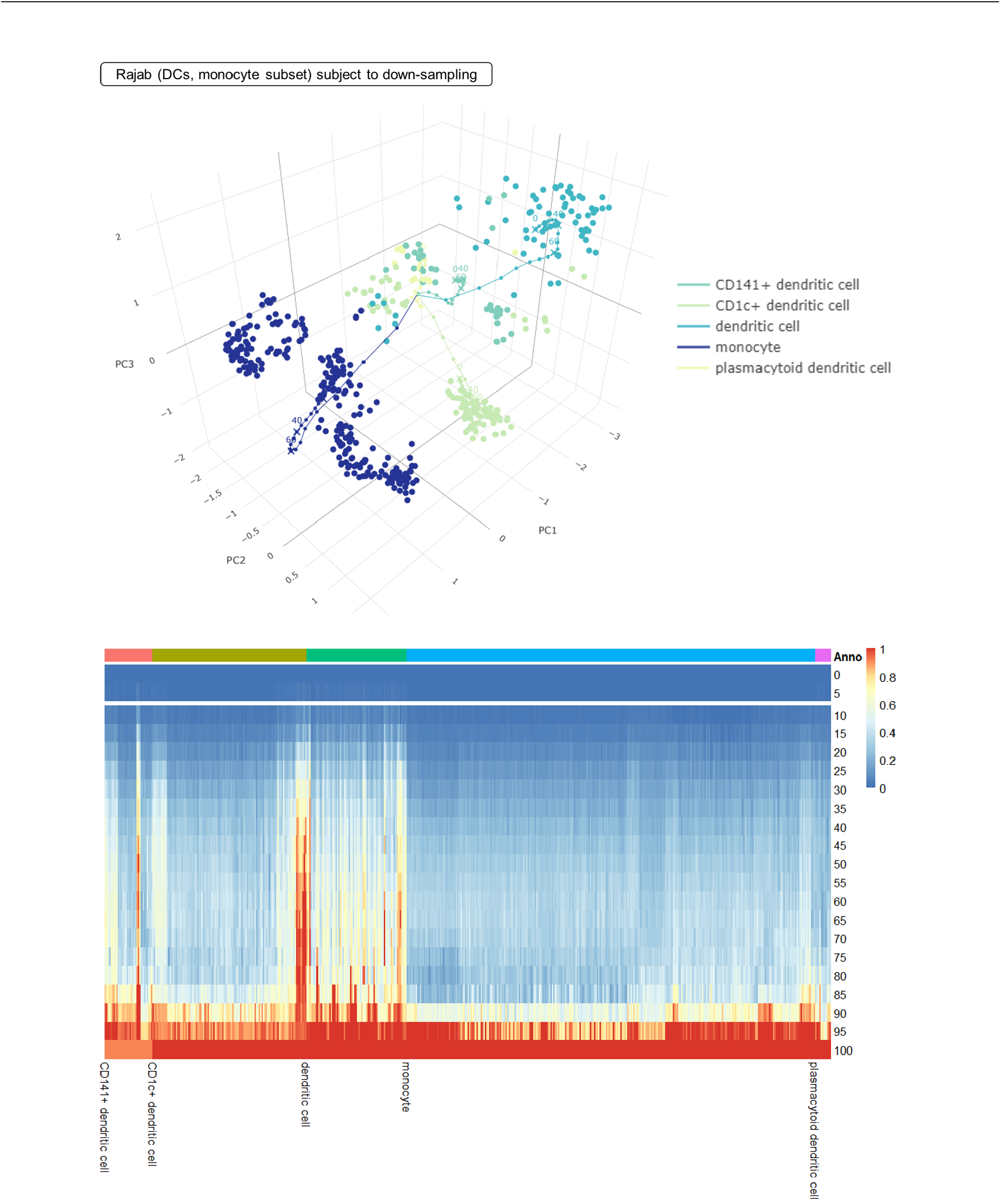
Impact of sparsity on projecting cells to the Monocyte-Dendritic cell subset of Rajab et al. (2021) atlas with an analysis similar to Figures S6, S7 and S8. When sparsity is less than 10 percent, 90 percent of pseudo-single cells retained distance ratio smaller than 0.5. We conclude that querying single cells on the Rajab et al. (2021) atlas may not be impacted when the sparsity of the query data is less than 10 percent.

**Figure S10.**
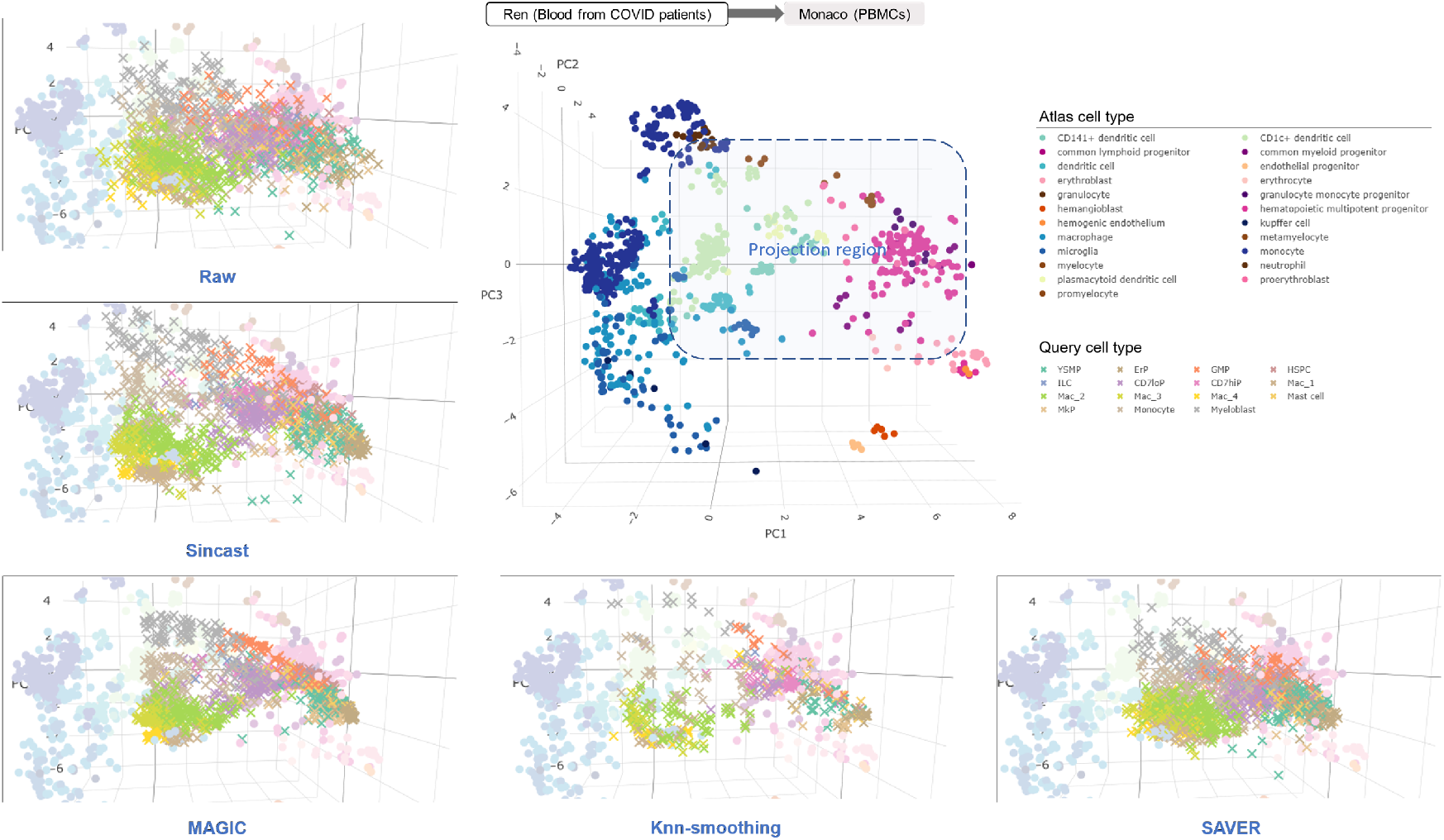
Projection of cells from Bian et al. (2020) on the Rajab et al. (2021) atlas when cell imputation is performed. Different imputation methods led to different data structures in the query due to their algorithmic assumptions. For example, the distribution of MAGIC and SAVER imputed data were shrunk towards the middle of the atlas, potentially indicating improper post-imputation data scaling. The distribution of knn-smoothing imputed data was scattered as a result of aggregating cells locally at each cell’s neighborhood.

**Figure S11.**
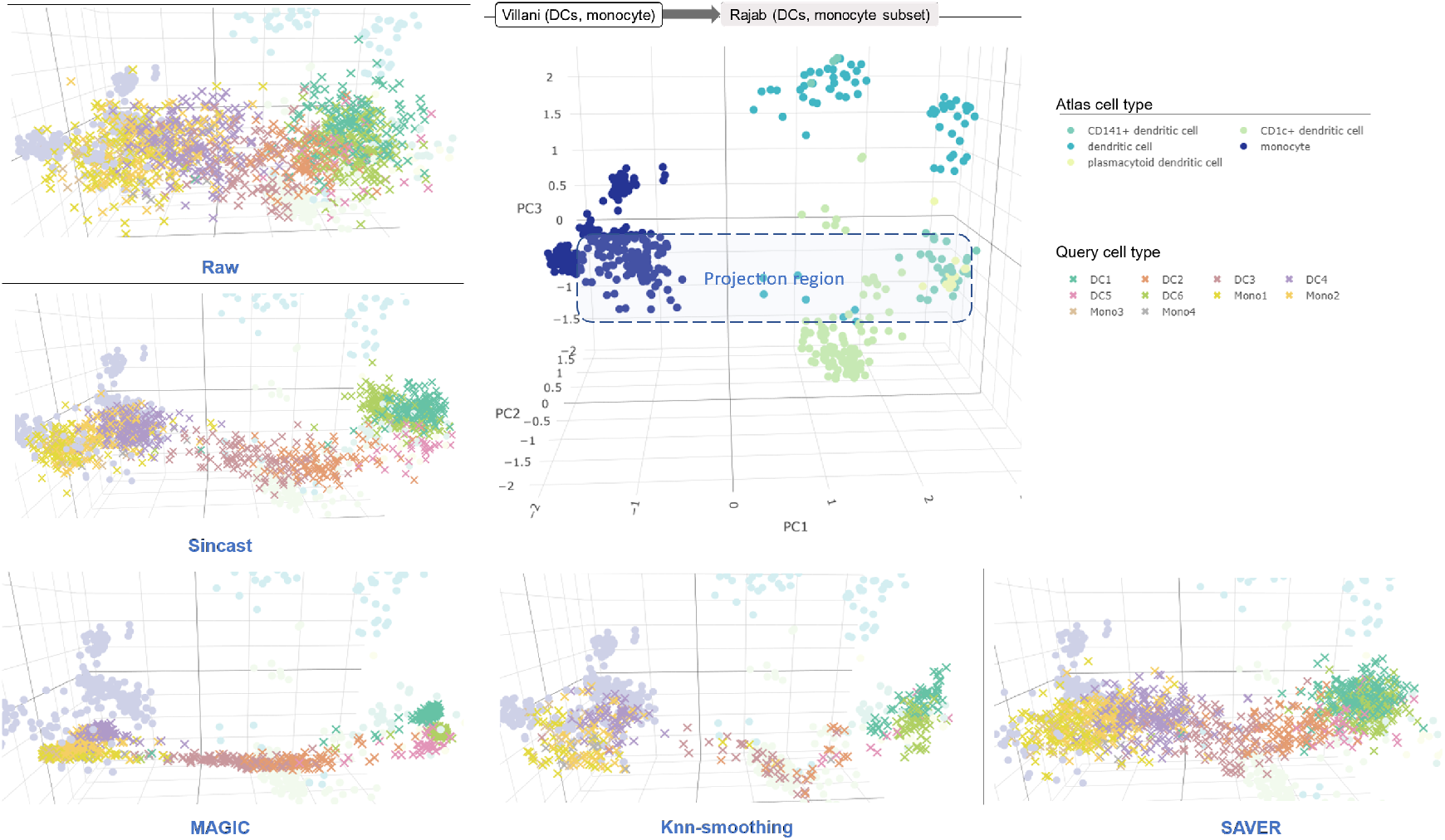
Projection of cells from Villani et al. (2017) imputed by different imputation methods, on the Monocyte-Dendritic cell subset of the Rajab et al. (2021) atlas.

**Figure S12.**
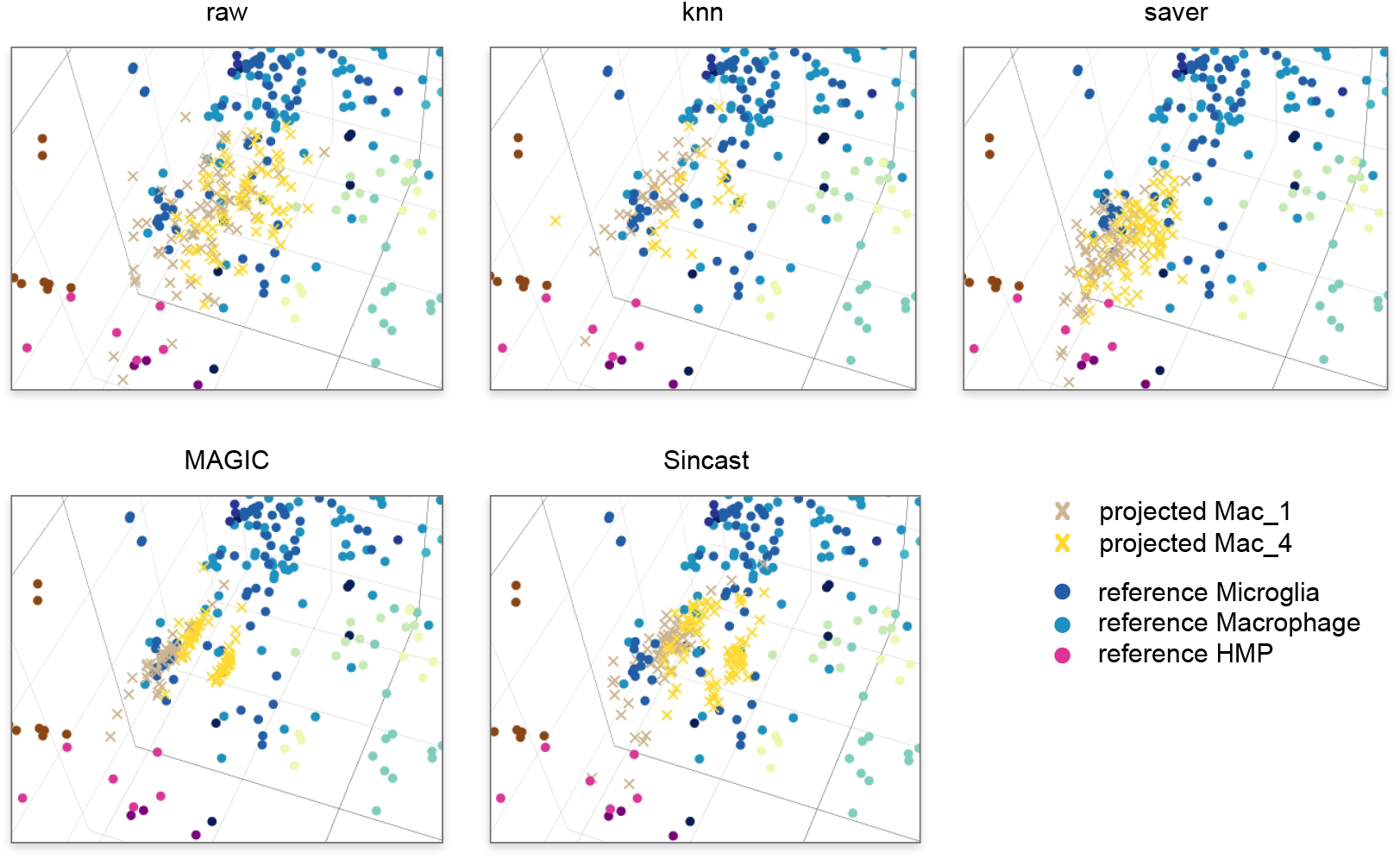
Mac_1_ and Mac_4_ query cells from Bian et al. (2020) were imputed using different methods prior to projection onto the Rajab et al. (2021) atlas. Close-up on the relevant area of PCA space is shown. Only MAGIC and Sincast imputation methods result in separation of these two clusters when projected, consistent with observations made by Bian et al. (2020).

**Figure S13.**
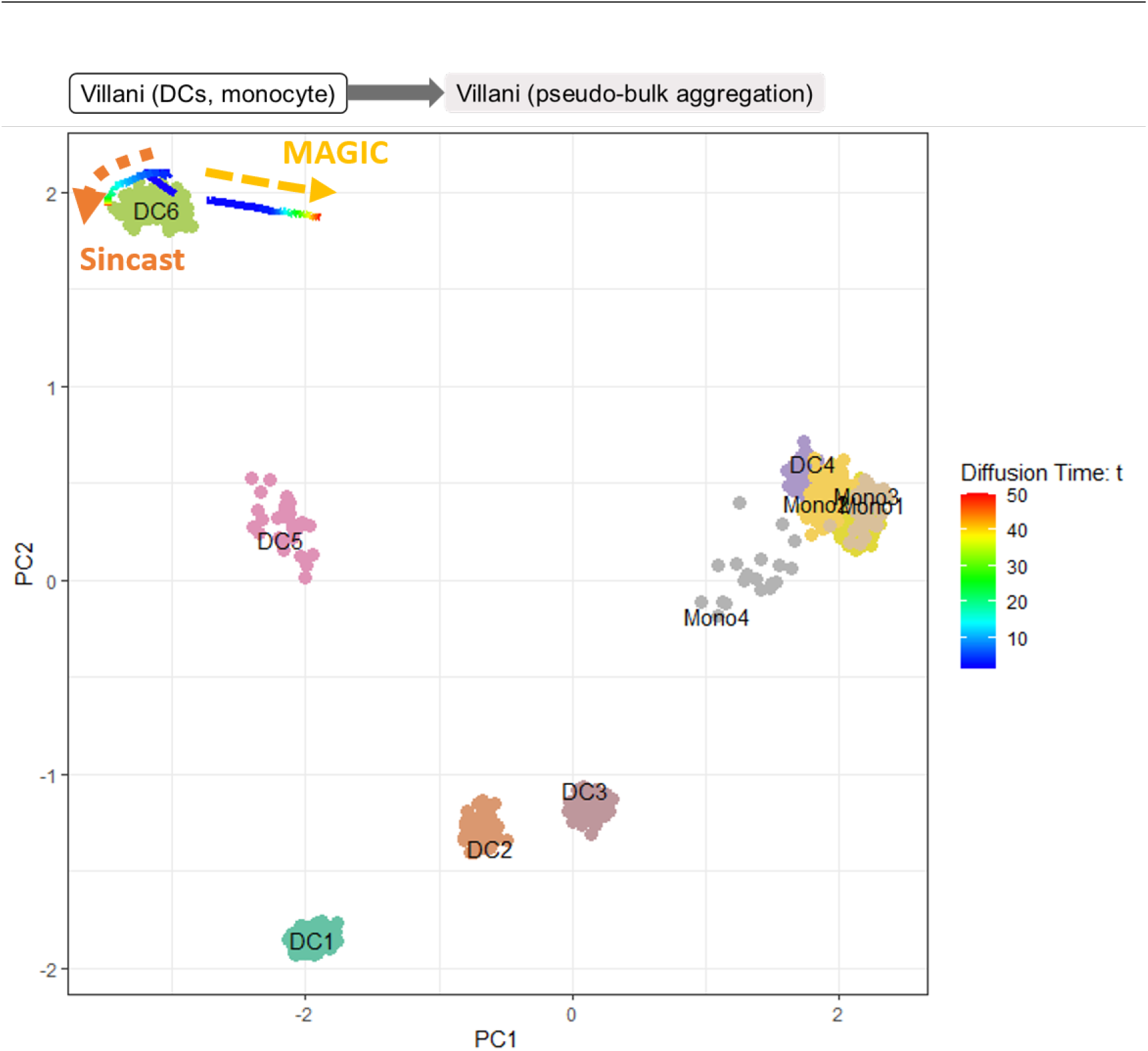
Projection of Sincast and the MAGIC imputed DC6 (pDC) cell population from Villani et al. (2017) onto the monocyte-DC subset of the Rajab et al. (2021) atlas. Imputation was performed on the 20 DC6 and 287 monocyte subsets of the Villani et al. (2017) data (the number of DC6 is increased from 10 to 20 compared to Figure 3B). The imputation neighborhood size is set to 15, which is smaller than the actual size of DC6 population included in the test data. Projection centroids of the imputed DC6 population with increasing diffusion time *t* are shown. The shift of MAGIC imputed DC6 population towards monocyte identity was not as strong as in Figure 3B, suggesting strong impact of tuning on MAGIC imputation.

**Figure S14.**
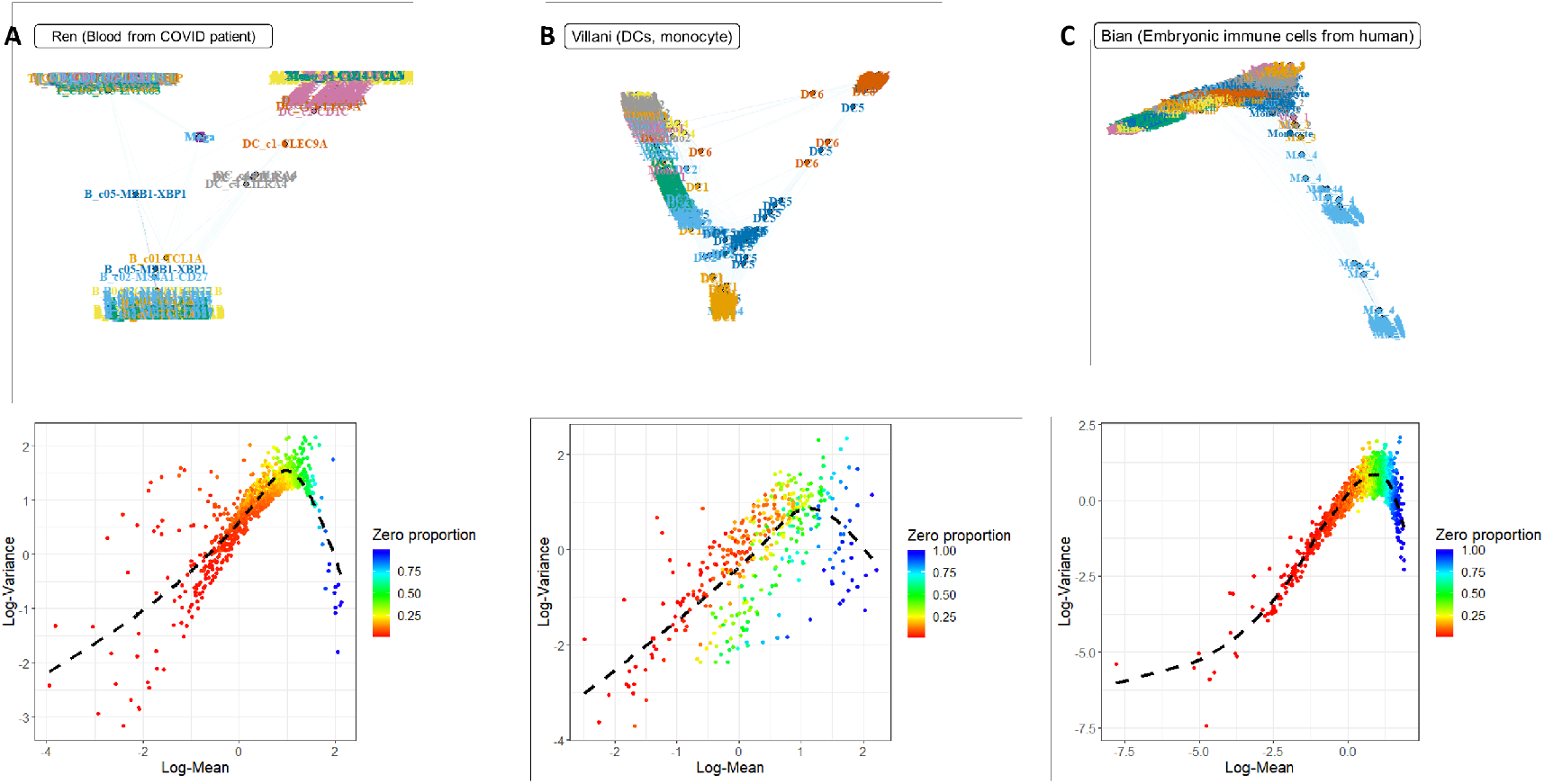
Diagnostic plots for Sincast imputation and post-imputation data scaling, made independently for **(A)** Ren et al. (2021), **(B)** Villani et al. (2017) and **(C)** Villani et al. (2017) data. For each query data, the top panel shows the diffusion embedding learnt by eigen decomposition of the diffusion operator used for Sincast data imputation. Cells in the embedding are connected by weighted lines (edges) representing affinities. The Sincast diffusion embedding gives some intuition on how the query cells are connected and hence impute each other in the graph defined by Sincast. For example, in (A) we can observe that the graph constructed was much sparser than in (B) and (C), informing that cells of different components in graph such as myeloid cells and lymphoid cells rarely impute each other. The bottom panel shows the log-gene mean and variance relationship representing global gene dispersion trend in the imputed data. Dashed black line represents generalized additive model fitting on the trend. Different data showed similar trend, suggesting that there are consistent dispersion trends to be estimated in scRNA-seq data. We scaled the imputed data according to their trend estimation.

**Figure S15.**
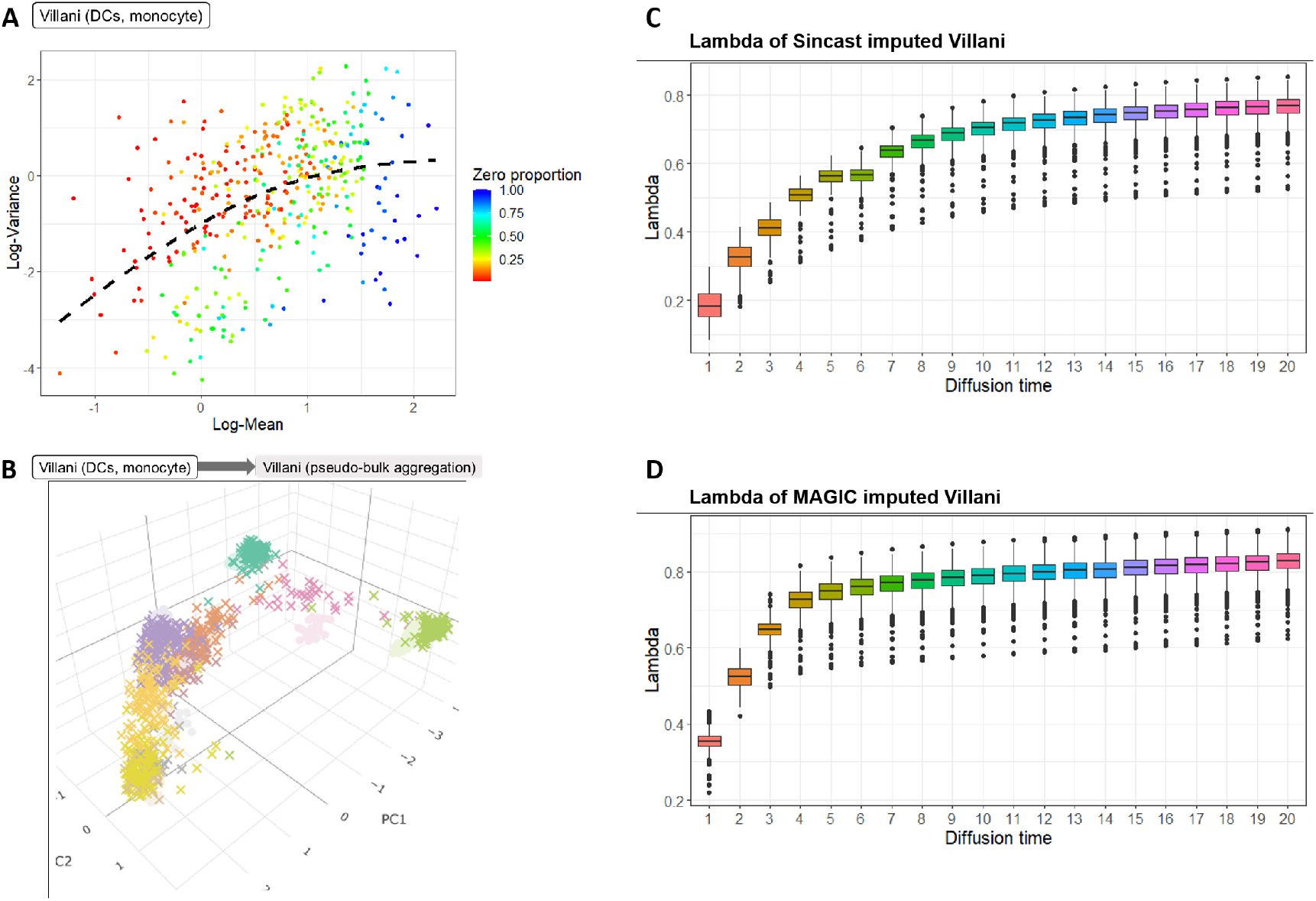
Sincast post-imputation data scaling on MAGIC imputed Villani et al. (2017) data. **(A)** Log-gene mean and variance relationship estimated on MAGIC imputed data (similar to Figure S14B, top panel for Sincast), showing that MAGIC and Sincast imputed data share similar dispersion trends. **(B)** Projection of MAGIC imputed-Sincast scaled data onto a pseudo-bulk version of the Villani et al. (2017) data. The loss of local and global variation after imputation was successfully recovered by Sincast data scaling (Compared to Figure 3A, yellow panel for MAGIC). **(C)** Distribution of scaling strength lambda for each imputed cell with Sincast. When lambda increases, the imputed cells tend to shrink back to the original, unimputed observations. As imputation strength increases (diffusion time *t*), lambda also increased in response. **(D)** Similar to (C) but for MAGIC imputed data. Compared to Sincast imputation, the lambda’s increase was more sharp as diffusion time increased, suggesting that MAGIC has a stronger imputation strength than Sincast, potentially leading to over-smoothing.

## References

1. Andreatta, M., Corria-Osorio, J., Müller, S., Cubas, R., Coukos, G., and Carmona, S. J. (2021). Interpretation of t cell states from single-cell transcriptomics data using reference atlases. Nature communications, 12(1):1–19.

2. Angel, P. W., Rajab, N., Deng, Y., Pacheco, C. M., Chen, T., Lê Cao, K.-A., Choi, J., and Wells, C. A. (2020). A simple, scalable approach to building a cross-platform transcriptome atlas. PLoS computational biology, 16(9):e1008219.

3. Aran, D., Looney, A. P., Liu, L., Wu, E., Fong, V., Hsu, A., Chak, S., Naikawadi, P., Wolters, P. J., Abate, A. R., et al. (2019). Reference-based analysis of lung single-cell sequencing reveals a transitional profibrotic macrophage. Nature immunology, 20(2):163–172.

4. Argelaguet, R., Cuomo, A. S., Stegle, O., and Marioni, J. C. (2021). Computational principles and challenges in single-cell data integration. Nature Biotechnology, pages 1–14.

5. Bian, Z., Gong, Y., Huang, T., Lee, C. Z., Bian, L., Bai, Z., Shi, H., Zeng, Y., Liu, C., He, J., et al. (2020). Deciphering human macrophage development at single-cell resolution. Nature, pages 1–6.

6. Bolstad, B. M., Irizarry, R. A., Åstrand, M., and Speed, T. P. (2003). A comparison of normalization methods for high density oligonucleotide array data based on variance and bias. Bioinformatics, 19(2):185–193.

7. Chandra, V., Bhattacharyya, S., Schmiedel, B. J., Madrigal, A., Gonzalez-Colin, C., Fotsing, S., Crinklaw, A., Seumois, G., Mohammadi, P., Kronenberg, M., et al. (2021). Promoter-interacting expression quantitative trait loci are enriched for functional genetic variants. Nature Genetics, 53(1):110–119.

8. Choi, J., Pacheco, C. M., Mosbergen, R., Korn, O., Chen, T., Nagpal, I., Englart, S., Angel, P. W., and Wells, C. A. (2019). Stemformatics: visualize and download curated stem cell data. Nucleic acids research, 47(D1):D841–D846.

9. Cieslak, D. A., Hoens, T. R., Chawla, N. V., and Kegelmeyer, W. P. (2012). Hellinger distance decision trees are robust and skew-insensitive. Data Mining and Knowledge Discovery, 24(1):136–158.

10. Clarke, Z. A., Andrews, T. S., Atif, J., Pouyabahar, D., Innes, B. T., MacParland, A., and Bader, G. D. (2021). Tutorial: guidelines for annotating single-cell transcriptomic maps using automated and manual methods. Nature protocols, 16(6):2749–2764.

11. Cobos, F. A., Alquicira-Hernandez, J., Powell, J. E., Mestdagh, P., and De Preter, K. (2020). Benchmarking of cell type deconvolution pipelines for transcriptomics data. Nature communications, 11(1):1–14.

12. Coifman, R. R. and Lafon, S. (2006). Diffusion maps. Applied and computational harmonic analysis, 21(1):5–30.

13. Crowell, H. L., Soneson, C., Germain, P.-L., Calini, D., Collin, L., Raposo, C., Malhotra, D., and Robinson, M. D. (2020). Muscat detects subpopulation-specific state transitions from multi-sample multi-condition single-cell transcriptomics data. Nature communications, 11(1):1–12.

14. Davis, C. A., Hitz, B. C., Sloan, C. A., Chan, E. T., Davidson, J. M., Gabdank, I., Hilton, J. A., Jain, K., Baymuradov, U. K., Narayanan, A. K., et al. (2018). The encyclopedia of dna elements (encode): data portal update. Nucleic acids research, 46(D1):D794–D801.

15. Fu, G.-H., Wu, Y.-J., Zong, M.-J., and Pan, J. (2020). Hellinger distance-based stable sparse feature selection for high-dimensional class-imbalanced data. BMC bioinformatics, 21(1):1–14.

16. Hao, Y., Hao, S., Andersen-Nissen, E., Mauck III, W. M., Zheng, S., Butler, A., Lee, M. J., Wilk, A. J., Darby, C., Zager, M., et al. (2021). Integrated analysis of multimodal single-cell data. Cell.

17. Hennig, C. (2020). fpc: Flexible Procedures for Clustering. R package version 2.2–5.

18. Hou, W., Ji, Z., Ji, H., and Hicks, S. C. (2020). A systematic evaluation of single-cell rna-sequencing imputation methods. Genome biology, 21(1):1–30.

19. Huang, M., Wang, J., Torre, E., Dueck, H., Shaffer, S., Bonasio, R., Murray, J. I., Raj, A., Li, M., and Zhang, N. R. (2018). Saver: gene expression recovery for single-cell rna sequencing. Nature methods, 15(7):539–542.

20. Hubert, L. and Arabie, P. (1985). Comparing partitions. Journal of classification, 2(1):193–218.

21. Kassambara, A. and Mundt, F. (2017). Package ‘factoextra’. Extract and visualize the results of multivariate data analyses, 76.

22. Kong, W., Fu, Y. C., and Morris, S. A. (2020). Capybara: A computational tool to measure cell identity and fate transitions. bioRxiv.

23. Kuksin, M., Morel, D., Aglave, M., Danlos, F.-X., Marabelle, A., Zinovyev, A., Gautheret, D., and Verlingue, L. (2021). Applications of single-cell and bulk rna sequencing in onco-immunology. European Journal of Cancer, 149:193–210.

24. Lizio, M., Harshbarger, J., Shimoji, H., Severin, J., Kasukawa, T., Sahin, S., Abugessaisa, I., Fukuda, S., Hori, F., Ishikawa-Kato, S., et al. (2015). Gateways to the fantom5 promoter level mammalian expression atlas. Genome biology, 16(1):1–14.

25. Love, M. I., Huber, W., and Anders, S. (2014). Moderated estimation of fold change and dispersion for rna-seq data with deseq2. Genome biology, 15(12):1–21.

26. Luecken, M. D., Büttner, M., Chaichoompu, K., Danese, A., Interlandi, M., Mueller, M. F., Strobl, D. C., Zappia, L., Dugas, M., Colomé-Tatché, M., et al. (2020). Benchmarking atlas-level data integration in single-cell genomics. BioRxiv.

27. Mabbott, N. A., Baillie, J. K., Brown, H., Freeman, T. C., and Hume, D. A. (2013). An expression atlas of human primary cells: inference of gene function from coexpression networks. BMC genomics, 14(1):1–13.

28. McInnes, L., Healy, J., and Melville, J. (2018). Umap: Uniform manifold approximation and projection for dimension reduction. arXiv preprint arXiv:1802.03426.

29. Monaco, G., Lee, B., Xu, W., Mustafah, S., Hwang, Y. Y., Carre, C., Burdin, N., Visan, L., Ceccarelli, M., Poidinger, M., et al. (2019). Rna-seq signatures normalized by mrna abundance allow absolute deconvolution of human immune cell types. Cell reports, 26(6):1627–1640.

30. Moon, K. R., van Dijk, D., Wang, Z., Gigante, S., Burkhardt, D. B., Chen, W. S., Yim, K., van den Elzen, A., Hirn, M. J., Coifman, R. R., et al. (2019). Visualizing structure and transitions in high-dimensional biological data. Nature biotechnology, 37(12):1482–1492.

31. Peng, T., Zhu, Q., Yin, P., and Tan, K. (2019). Scrabble: single-cell rna-seq imputation constrained by bulk rna-seq data. Genome biology, 20(1):88.

32. Petković, M., Škrlj, B., Kocev, D., and Simidjievski, N. (2020). Fuzzy jaccard index: A robust comparison of ordered lists. arXiv preprint arXiv:2008.02216.

33. Picelli, S., Björklund, Å. K., Faridani, O. R., Sagasser, S., Winberg, G., and Sandberg, R. (2013). Smart-seq2 for sensitive full-length transcriptome profiling in single cells. Nature methods, 10(11):1096–1098.

34. Rajab, N., Angel, P. W., Deng, Y., Gu, J., Jameson, V., Kurowska-Stolarska, M., Milling, S., Pacheco, C. M., Rutar, M., Laslett, A. L., et al. (2021). An integrated analysis of human myeloid cells identifies gaps in in vitro models of in vivo biology. Stem cell reports, 16(6):1629–1643.

35. Ren, X., Wen, W., Fan, X., Hou, W., Su, B., Cai, P., Li, J., Liu, Y., Tang, F., Zhang, F., et al. (2021). Covid-19 immune features revealed by a large-scale single-cell transcriptome atlas. Cell, 184(7):1895–1913.

36. Richards, J. and Cannoodt, R. (2019). diffusionMap: Diffusion Map. R package version 1.2.0.

37. Roels, J., Kuchmiy, A., De Decker, M., Strubbe, S., Lavaert, M., Liang, K. L., Leclercq, G., Vandekerckhove, B., Van Nieuwerburgh, F., Van Vlierberghe, P., et al. (2020). Distinct and temporary-restricted epigenetic mechanisms regulate human *a/3* and *y8* t cell development. Nature Immunology, 21(10):1280–1292.

38. Rousseeuw, P. J. (1987). Silhouettes: a graphical aid to the interpretation and validation of cluster analysis. Journal of computational and applied mathematics, 20:53–65.

39. Sarkar, A. and Stephens, M. (2021). Separating measurement and expression models clarifies confusion in single-cell rna sequencing analysis. Nature Genetics, 53(6):770–777.

40. Schmiedel, B. J., Singh, D., Madrigal, A., Valdovino-Gonzalez, A. G., White, B. M., Zapardiel-Gonzalo, J., Ha, B., Altay, G., Greenbaum, J. A., McVicker, G., et al. (2018). Impact of genetic polymorphisms on human immune cell gene expression. Cell, 175(6):1701–1715.

41. Street, K., Risso, D., Fletcher, R. B., Das, D., Ngai, J., Yosef, N., Purdom, E., and Dudoit, S. (2018). Slingshot: cell lineage and pseudotime inference for single-cell transcriptomics. BMC genomics, 19(1):1–16.

42. Tang, K., Ji, X., Zhou, M., Deng, Z., Huang, Y., Zheng, G., and Cao, Z. (2021). Rank-in: enabling integrative analysis across microarray and rna-seq for cancer. Nucleic Acids Research.

43. Thul, P. J., Åkesson, L., Wiking, M., Mahdessian, D., Geladaki, A., Blal, H. A., Alm, T., Asplund, A., Björk, L., Breckels, L. M., et al. (2017). A subcellular map of the human proteome. Science, 356(6340).

44. Van Dijk, D., Sharma, R., Nainys, J., Yim, K., Kathail, P., Carr, A. J., Burdziak, C., Moon, K. R., Chaffer, C. L., Pattabiraman, D., et al. (2018). Recovering gene interactions from single-cell data using data diffusion. Cell, 174(3):716–729.

45. Villani, A.-C., Satija, R., Reynolds, G., Sarkizova, S., Shekhar, K., Fletcher, J., Griesbeck, M., Butler, A., Zheng, S., Lazo, S., et al. (2017). Single-cell rna-seq reveals new types of human blood dendritic cells, monocytes, and progenitors. Science, 356(6335).

46. Wagner, F., Yan, Y., and Yanai, I. (2017). K-nearest neighbor smoothing for high-throughput single-cell rna-seq data. BioRxiv, page 217737.

47. Wood, S. N. (2011). Fast stable restricted maximum likelihood and marginal likelihood estimation of semiparametric generalized linear models. Journal of the Royal Statistical Society: Series B (Statistical Methodology), 73(1):3–36.

48. Yip, S. H., Sham, P. C., and Wang, J. (2019). Evaluation of tools for highly variable gene discovery from single-cell rna-seq data. Briefings in bioinformatics, 20(4):1583–1589.

49. Zhang, D., Guo, R., Lei, L., Liu, H., Wang, Y., Wang, Y., Qian, H., Dai, T., Zhang, T., Lai, Y., et al. (2020). Covid-19 infection induces readily detectable morphologic and inflammation-related phenotypic changes in peripheral blood monocytes. Journal of leukocyte biology.

50. Zhao, X., Wu, S., Fang, N., Sun, X., and Fan, J. (2020). Evaluation of single-cell classifiers for single-cell rna sequencing data sets. Briefings in bioinformatics, 21(5):1581–1595.

51. Zheng, G. X., Terry, J. M., Belgrader, P., Ryvkin, P., Bent, Z. W., Wilson, R., Ziraldo, S. B., Wheeler, T. D., McDermott, G. P., Zhu, J., et al. (2017). Massively parallel digital transcriptional profiling of single cells. Nature communications, 8(1):1–12.

52. Zhou, Y., Fu, B., Zheng, X., Wang, D., Zhao, C., Qi, Y., Sun, R., Tian, Z., Xu, X., and Wei, H. (2020). Pathogenic t-cells and inflammatory monocytes incite inflammatory storms in severe covid-19 patients. National Science Review, 7(6):998–1002.

